# HIV-1 Ribonucleoprotein Complex Organization Facilitates Assembly of the Mature HIV-1 Capsid

**DOI:** 10.64898/2026.10.05.756791

**Authors:** Curt Waltmann, Manish Gupta, Alan N. Engelman, Dmitry Lyumkis, Gregory A. Voth

## Abstract

The encapsulation of the viral RNA condensate inside the capsid is a crucial step in the late stages of the HIV-1 replication cycle. While it is agreed that RNA encapsulation depends on RNA-bound integrase (IN), that capsid-bound IN-RNA filaments are present within the capsid, and that the ribonucleoprotein (RNP) complex takes up only a fraction of the capsid volume, the timing of filament formation and ribonucleoprotein (RNP) encapsulation are unclear. Here, we ran virion-scale, coarse grained molecular dynamics simulations to identify how the organization of the RNP and IN in particular affects mature capsid assembly and genome encapsulation. These virion-scale simulations allow for direct comparison between experimental capsid morphologies and those self-assembled in the simulations. It was found, when relatively few tetramers are on the RNP surface, that IN tetramers could facilitate RNA encapsulation without filling a significant fraction of the capsid volume, but do not form a capsid-templated filament. When octameric IN-RNA filaments are modeled, it was found that WT-like capsid morphologies form only when the IN is organized as a single long filament, in agreement with recent experimental suggestions. These capsids are more conical and elongated than the capsids assembled around the RNP with tetrameric IN. Thus, we suggest the IN-RNA filament assembles ahead of the capsid and directs it to its final conical morphology, consistent with the experimental observation of less elongated capsids when IN oligomerization is disrupted.

## Introduction

The HIV-1 capsid is a multipurpose container that secures the viral RNA (vRNA) genome from cellular immune sensors during cytoplasmic transport(1), acts as a bioreactor for early reverse transcription(2), enables nuclear import of the pre-integration complex (PIC) (3, 4), and releases the PIC into the host nucleus(5). A typical mature capsid contains more than 1200 CA proteins arranged in pentameric and hexameric organizations, creating a fullerene geometry(6, 7). While multiple *in vitro* assembly studies have demonstrated that a minimal combination of CA protein and the small anion inositol hexakisphosphate (IP6) is adequate for achieving the conical morphology(8-11), the capsid must also package the ribonucleoprotein (RNP) complex composed of the vRNA, nucleocapsid (NC) protein, integrase (IN) and reverse transcriptase to produce an infectious virion(12, 13). Packaging failure during maturation leads to the degradation of the unprotected viral genetic material by host defense systems(14). Interestingly, the RNP condensate has also been implicated in influencing CA assembly kinetics during biochemical reconstitutions of HIV-1 capsid maturation(15, 16). Thus, the mechanistic details of simultaneous capsid assembly and RNP packaging are critically important to understanding the capsid-RNP complex.

During maturation, protease (PR) mediated cleavage of the immature Gag lattice frees the subdomains of Gag from the Gag polyprotein, allowing the complete reconfiguration of the immature virion(17, 18). This includes ∼3000 copies of the CA protein, of about half of which will reassemble into the mature capsid(19, 20). Individual CA proteins contain a C-terminal and N-terminal domain (CTD and NTD) connected by a flexible linker(21). In solution, these proteins exist in a dynamic equilibrium between monomeric and dimeric states(22). Dimers are formed via CTD-CTD contacts, while contacts within hexamers and pentamers are formed by NTD-NTD contacts(23-25). The small anion cofactor IP6 stabilizes the pentamers(26, 27) which are needed to form the fullerene lattice(9, 11). Previous work has shown that IP6 binds at the inner pore of CA hexamers and pentamers via electrostatic interactions, leading to conical shapes(10) in the absence of the RNP. 3D imaging of radiation-sensitive NC by cryo-electron tomography and conventional electron microscopy of plastic-embedded thin sections showed that the RNP complex is predominantly located at the wide end of the cone(28), consistent with recent experiments that predicted that the RNP occupies ∼75% of the volume of the capsid core (29). This behavior is critical for the viral replication cycle as, during reverse transcription, the capsid interior must contain sufficient space to allow for the production of dsDNA(30).

*In vitro* approaches have identified several important features of the RNP complex (31, 32). Positively charged NC proteins are thought to play a key role in condensing the RNA dimer, which can be quite large due to self-repulsion brought on by its high negative charge. It is not known if all the 3000 available NC proteins are a part of the final RNP complex. However, just two thirds would contribute +22000 total charges and would be more than enough to compensate the -18000 total negative charges found on the two 9 kb strands of viral RNA(33, 34). This condensation mechanism, termed polyelectrolyte complexation(35-37) in some contexts, is ubiquitous in biomolecular condensates(38, 39).

The primary role of IN is to catalyze the integration of viral dsDNA into host chromatin(40). However, early viral genetic research also revealed pleiotropic outcomes of IN mutagenesis, including defects in particle assembly and release from producer cells(41). These were subsequently termed “class II mutations”. Notably, viruses harboring class II IN mutations produce increased proportions of empty capsids with mislocalized RNPs, pointing to a role for IN in assisting RNP localization in the late stages of the replication cycle (28). Further insight into the class II phenotype came from more recent observations showing how IN can directly bind to viral RNA to promote RNP condensation(42). Minimally, the IN tetramer is necessary for RNA binding and RNP condensate formation(43, 44). The structural basis of IN-RNA interactions was recently elucidated (45). IN forms octameric RNA filaments composed of interlinked tetramers that further interact with the capsid interior. These observations raise several fundamental questions about the requirements for a “packageable” RNP and the functional role of IN. What are the basic physical requirements for a packageable RNP? Is IN-RNA filament formation or higher-order assembly beyond tetramers necessary to fulfill these requirements? Does the capsid direct filament assembly or does the filament direct capsid assembly?

Large-scale all-atom molecular dynamics (AAMD) simulations have been employed to investigate the physical properties of the capsid-RNP complex(24, 45). However, the spatial and temporal limitations of AA models prevent them from simulating the dynamics of RNP encapsulation during capsid assembly Considering this, coarse-grained molecular dynamics(46) (CGMD) is the preferred approach to studying processes such as the assembly of HIV-1(4, 10, 47-52). Several previous phenomenological CG models have shown that genomes of many viruses are ideally matched to the size of their capsids(53) and thus do not describe the behavior of the partially empty HIV-1 capsid-RNP complex. This is also the case when the capsid itself is pleomorphic(54, 55) as is the case for HIV-1. One exception showed that partially empty shells can form when multiple types of shell proteins are present(56); however, this is also not the case for the mature HIV-1 capsid-RNP complex. The Voth group has created CGMD models to specifically study the assembly of the HIV-1 capsid(50, 51, 57) in the presence of host factor IP6(10), small molecule drug PF74 or Lenacapavir(48, 51), and have provided key insights into viral morphogenesis, despite the lack of an RNP model.

In this study, we extended the prior self-assembling CG capsid-IP6 model by including multiple RNP models. We used these models to investigate the underlying mechanisms that enable RNP enclosure within a capsid. We find that an RNP model with uniformly distributed capsid interactions was unable to reproduce the partially empty wild-type capsid structure, regardless of its physical properties and interaction strength. By adding a small domain of the RNP with different interactions we resolved this issue, implying that the multicomponent nature of the RNP is critical to assemble partially empty capsids. Next, we performed assembly simulations with a previously reported RNP model(12) updated using structural data for the IN tetramer (43) and parameterized for CGMD simulation. Using this integrative RNP model, we observed successful encapsulation when there were a few IN tetramers on the surface of the RNP condensate globule, with smaller capsids produced as more IN tetramers were located on the surface. Interestingly, when we allowed the IN tetramers more mobility within the RNP, we observed that clusters of IN tetramers formed without any attractions between the tetramers, suggesting that the capsid could induce IN tetramer clustering, but long, filament-like structures were not observed.. Importantly, when we simulated the assembly around pre-assembled octameric IN-RNA filaments(45), we found that having one long filament was preferred over three or more shorter ones, in agreement with experiments and the above observation that modeling multiple tetramers on the RNP surface yielded smaller capsids. These capsids were more elongated and conical than those assembled with IN tetramers, which we conclude agrees very well with experimental capsid ensembles. Taken together, our extensive capsid-RNP simulations support a preferred mechanism in which one long IN-RNA filament directs capsid assembly towards elongated conical morphologies.

## Results

We utilized a previously developed self-assembling CG model of a CA dimer(10, 50, 57), which is shown in Figure 1A. The CA proteins assemble into hexamers and pentamers shown in Figure 1B. As in previous work(10), our model includes IP6, which binds both the hexamer and the pentamer, and promotes the formation of conical capsids (11, 27). In our model, a fraction of the CA monomers adopt the assembly competent, “active” state and interact with other “active” CA monomers. Experiments show that in solution only ∼ 5-10% of CA monomers are in conformations where assembly is possible(22). We used the active fraction of CA dimers, [CA]_+_=10%, unless otherwise noted. Here, we developed three different RNP models (Figure 1D(i,ii,iii)), which we used throughout the manuscript. These RNP models include the effects of NC, which condenses the RNP implicitly, while Figure 1Dii and iii include the CG models of IN tetramers and filaments, respectively (Figure 1C). More details on the RNP models are included in the relevant sections of the text and the Methods Section. Figure 1E shows that our simulations also feature “crowders”, which are volume-excluding spheres that help to replicate the dense, crowded environment of the virion. We simulated systems of two different sizes. In the first case (Figure 1F(iii)), we simulate 750 CA dimers, 2128 crowders (to mimic the virion crowded interior)(50), 521 IP6 anions, and one RNP model in an 88 nm cubic box with periodic boundary conditions. The other case is meant to represent the stoichiometry and boundary conditions of the real virion with 1500 CA dimers, 4256 crowders, 1042 IP6 anions, and an explicit RNP model in a 140 nm diameter spherical confinement (to represent the virion envelope) made of immobile particles, as shown in Figure 1E(i,ii). These full virion-scale assembly simulations allowed us to assemble capsids directly comparable in size to those seen in virus-based experiments(13, 23). In both cases, we could assemble capsid-RNP complexes (Figure 1F).

**Figure 1.**
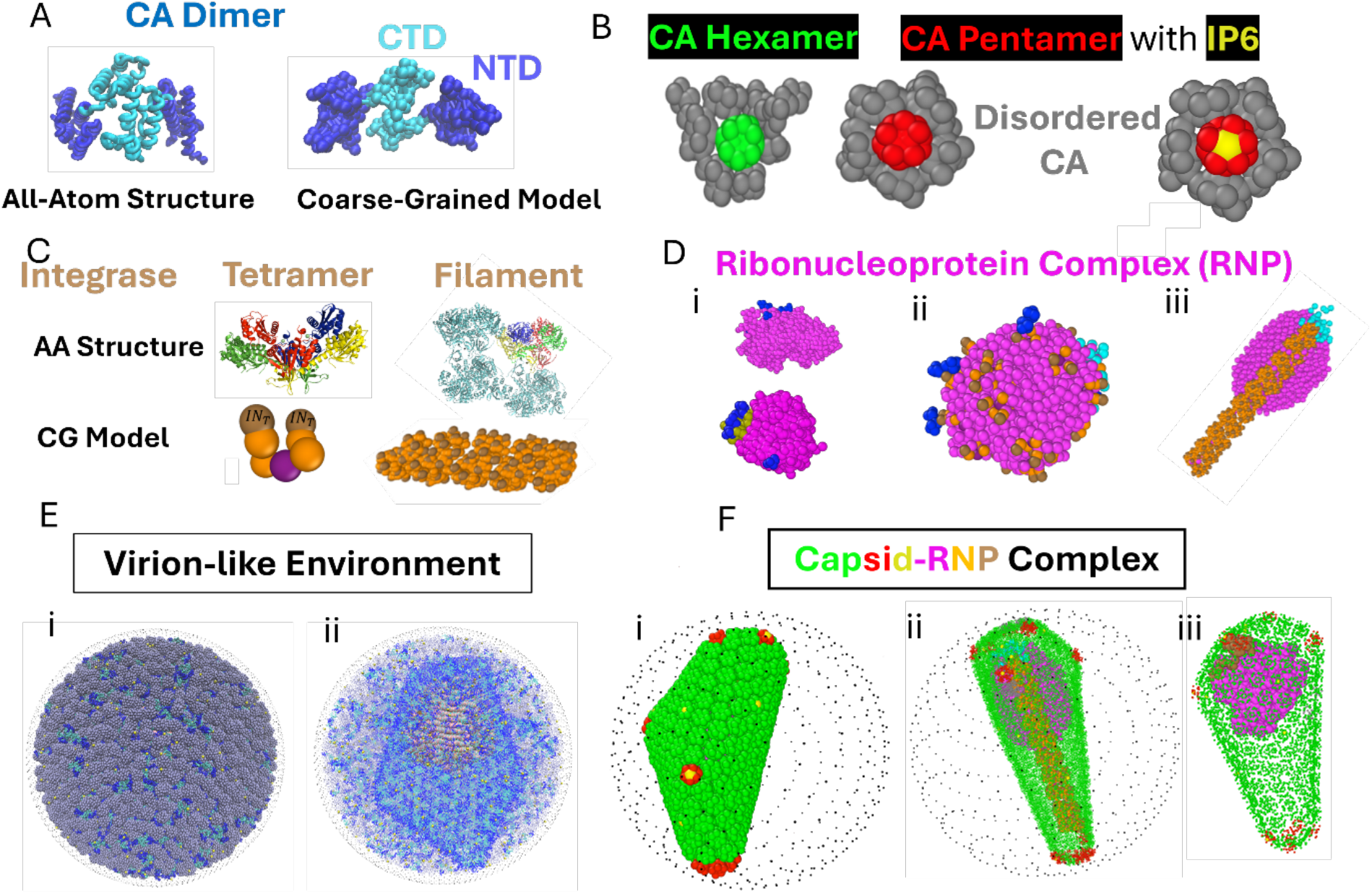
CG Model of Mature Capsid-RNP Assembly. (A) The smallest unit of assembly in our model is the CA dimer, which is comprised of two CA monomers, which feature a C-terminal domain (light blue) and an N-terminal domain (dark blue). The CG model of the CA dimer is based on a hetero elastic network model that captures the active conformations of the all-atom structure. (B) Many of these dimers combine to form CA Hexamers (green) and CA pentamers (red). Disordered CA (gray) is found at the edges of these growing assemblies. Hexamers and pentamers can both bind the small anion, inositol hexakisophosphate (IP6) (yellow), although there is a preference for pentamers. (C) All-atom and CG models of the IN tetramer and filament, which interact with the CA through the tan CG beads found on each structure. All-atom chains colored red, blue, yellow, and green show the tetramer within the filament(45) are quite similar to the individual tetramers(43), save for the configuration of the CTDs. (D) Tetramers and filament models are incorporated into the models of the RNP(ii and iii respectively), while the interactions with the capsid from the RNPs shown in (i) are found in the pink domains all over or in the small tan domains, which represent the RNA packaging signal. All models of the RNP implicitly contain the nucleocapsid which condenses the RNA and have a cyan region which is the dimerized 5’ untranslated region (UTR)(58). (E) Assembly occurs in a dense environment featuring inert crowder molecules (gray spheres) and either a repeating boundary condition or a membrane-like sphere (black dots) that simulates the membrane of the virion. (F) All of these components combine to form the mature capsid-RNP complex as shown. CA proteins made more transparent (in ii and iii) such that the RNP can be visualized within the capsid.

Two types of CGMD simulations were run. The first were nucleation simulations where CA dimers were run for up to 30 million CG timesteps and then stopped to look for a nucleus comprised of at least one hexamer or pentamer. (It is to be noted that CGMD time, by methodological design, is greatly accelerated over real laboratory time so as to enhance the sampling.) These simulations helped to identify whether nucleation will occur in the bulk solution or on the RNP for the given set of parameters. The second type was the assembly simulation. Here, we started with a nucleated hexamer, which is the assembly seed. CA dimers that are a part of this seed were fixed in the active state, along with some fraction of free dimers outside the nucleating lattice. The active free dimers were updated every 5x10^5^ CGMD timesteps.

### One component RNP models assemble into capsid-RNP complexes with improper structure

To understand how RNP organization impacts its properties relative to capsid assembly, we began our investigation by considering all components of the RNP to be uniformly distributed, lac king tan packaging domains (Figure 1D(i)). Thus, under these conditions, all interactions between CA proteins and the RNP can be thought of as the average of the interactions between CA proteins and each individual component of the RNP. The RNP was modeled as a one component polymer. We varied its length and condensation state (which is thought to be controlled by NC proteins) as well as its interactions with CA through the parameter ε_CA_ as shown in Figure 2A. To minimize computational costs, we used the smaller, 750 CA dimer system for these simulations. More details on the model can be found in the Methods section. In the simulations, we observed 4 different classes of assemblies as shown in Figure 2A, using an active CA assembly fraction of 15%. These classes were based on the length (N) and condensation state (X) of the RNP as well as the interaction with the CA protein (ε_CA_). When the interaction between the RNP and CA was too weak, we observed the formation of empty capsids, with the RNP mislocalized on the outside. This is shown in the left-most column of Figure 2 (red outline), where one can also see how the conformation of the RNP polymer changes based on the length and condensation state. The condensation state can be 1, 2, or 3, with 1 being the least condensed and 3 being the most condensed. Full details on how the condensation state is controlled can be found in the Supplementary Methods RNP Model 1 section. Empty capsids with external RNPs like these have been observed under a variety of conditions, including class II IN mutations(41, 42, 44, 59) and/or treatment with allosteric IN inhibitors(28, 42, 60, 61), or sublethal doses of protease inhibitors(62). When ε_CA_ was 1.2 kcal/mol and the RNP was not strongly condensed (X=1,2), we observed disordered assemblies. These tended to be malformed and unclosed and in the one case of closure, 20 pentamers as well as a variety of other defects were present. By strongly condensing the RNP (X=3), we observed encapsulation of the RNP. However, the resultant capsid shapes were highly spherical and non-canonical (see also Figure S1). This is reminiscent of viral capsids formed in the presence of a capsid inhibitor that binds the underside of hexamers(63) and thus may be affecting CA-RNP interactions. The size of these capsids was determined by the size of the RNP, which completely filled the capsid. By using an intermediate value of ε_CA_, such as 0.6 kcal/mol, and a weakly condensed RNP (X=1,2), we observed canonical capsid shapes that encapsulate the RNP. However, as in the case of the spherical capsids, these capsids were completely filled by the RNP. More examples are shown in Figure 2B. Although the genome of many viruses does fill the capsid and even determines the capsid size(53, 64), recent experiments have predicted that the RNP occupies ∼75% of the capsid volume (29). These observations suggest that a one-component mean field model of the RNP fails to recapitulate biological assembly of an optimal capsid-RNP complex.

**Figure 2.**
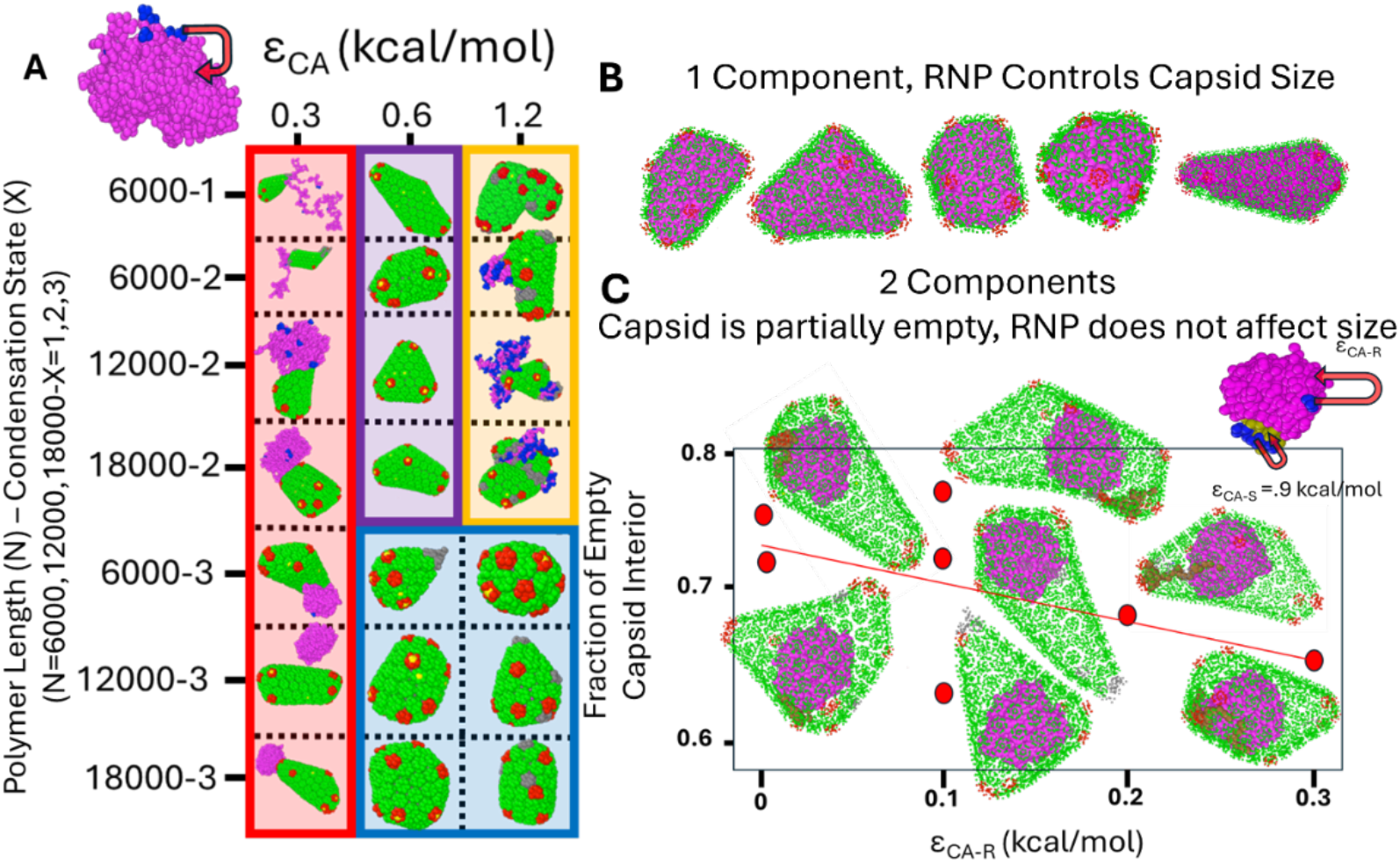
Capsid assembly around a simple 1 or 2 component RNP polymer. (A) Four different general classes of assembly were observed based on the physical parameters of the polymeric RNP model (polymer length and condensation state on the y axis) and the interaction between CA and the RNP, ε_CA_, on the x axis. These were empty capsids(red boxes), disordered assemblies(yellow boxes), spherical RNP-containing capsids(blue boxes), and canonical RNP-containing capsids(purple boxes). (B) additional replicates are shown with length 18000 and ε_CA_ =0.6 kcal/mol, where completely full canonical capsids formed. (C) The fraction of empty capsid interior is shown as a function of ε_CA-R_, the interaction between and the rest of the RNP, for 12000-monomer long RNPs. The CA interaction with the packaging domain/ signal was a constant 0.9 kcal/mol (top right). Snapshots of the capsid-RNP complex are visualized to emphasize the partially empty space within the capsid interior.

Since the one component model seemingly failed to construct biologically-relevant capsids, we hypothesized that a more complex model must be necessary. We added a second tan domain to our polymeric model of the RNP as shown in the top right of Figure 2C. This second tan domain could function as a packaging signal, like the role of psi RNA during the assembly of immature HIV-1 particles(65). This second domain was assigned a stronger interaction with CA (ε_CA-S_=0.9 kcal/mol), while the interaction of CA with the rest of the RNP, ε_CA-R_, was too weak to lead to attachment of the capsid. More details on the model can be found in the Supplementary Methods RNP Model 2 section. Thus, the second domain effectively served as the packaging signal and provided the necessary interactions between CA proteins and the RNP. More illustration of the kinetics of this process can be seen in Figure S2. Here, we only considered strongly condensed RNPs and polymer lengths of 12000 CG particles, which was one length used in experiments exploring how genome length affects uncoating(29). Figure 2C shows capsid-RNP complexes with only approximately 30% of the interior volume of the capsid filled. We note that larger RNPs in SI Figure S1 take up 40-70%. Thus, using smaller RNPs leads to more empty space in the capsid-RNP complex. This is consistent with recent experimental data showing that smaller RNPs do not efficiently uncoat, likely due to their inability to apply sufficient pressure on the capsid after reverse transcription(29).

### RNPs with IN also depend on surface concentration

The previous simulations showed that assembly of a partially empty capsid required a component interacting strongly with the capsid, while the rest of the RNP had little to no interaction. Recent experimental structures suggest that interacting component is the IN-RNA filament, while the rest of the RNP does not bind the capsid. These structures do not tell us about the order in which the IN-RNA filament and capsid assemble. Thus, we wondered if the RNP would be packageable before formation of the IN-RNA filament is complete. To test this, we explicitly included IN(42, 44) in the RNP as tetramers, which is the form required for RNA binding(43, 44). Although there are no experimentally-derived structures for IN tetramers bound to RNA, a previous modeling study(12) provided a hypothesis for the RNP structure, including the distribution of IN and structured regions of the vRNA. We also updated the IN model based on structural data for the apo IN tetramer, which was determined in the absence of bound nucleic acid(43) (see Figure 1C, D(ii) and the Supplementary Information). In this scenario, we used hypothetical interactions between CA and the RNA-bound IN tetramers (Figure S11) through the *IN*_*T*_sites (Figure 1C), since it is known the RNP is not packageable without IN(42, 44). We also used the virion-like assembly conditions (see Figure 1E) here and throughout the rest of the manuscript.

As shown in Figure 3, the new RNP model still has one free parameter, H_IN-R_, which is the strength of the interaction between the IN and the rest of the RNP (see Supplementary Methods RNP Model 3 for details). This parameter allowed us to control how much IN partitioned to the surface of the RNP and was thus available to make the hypothetical interaction with the capsid. In Figures 3A and B, we show two examples where the parameter is weak (0 kcal/mol) or strong (0.72 kcal/mol). In both cases, we quantified the degree of burial (DOB) for the brown IN *IN*_*T*_CG beads (the 2 beads at the ‘top’ of each tetramer, which interact with CA, see Figure S11 for more details), which is also known as the circular variance(66). It is defined below as 1 minus the vector sum of all unit vectors between an IN *IN*_*T*_bead and its RNP neighbors within a cut-off distance, *R*_*cut*_

**Figure 3.**
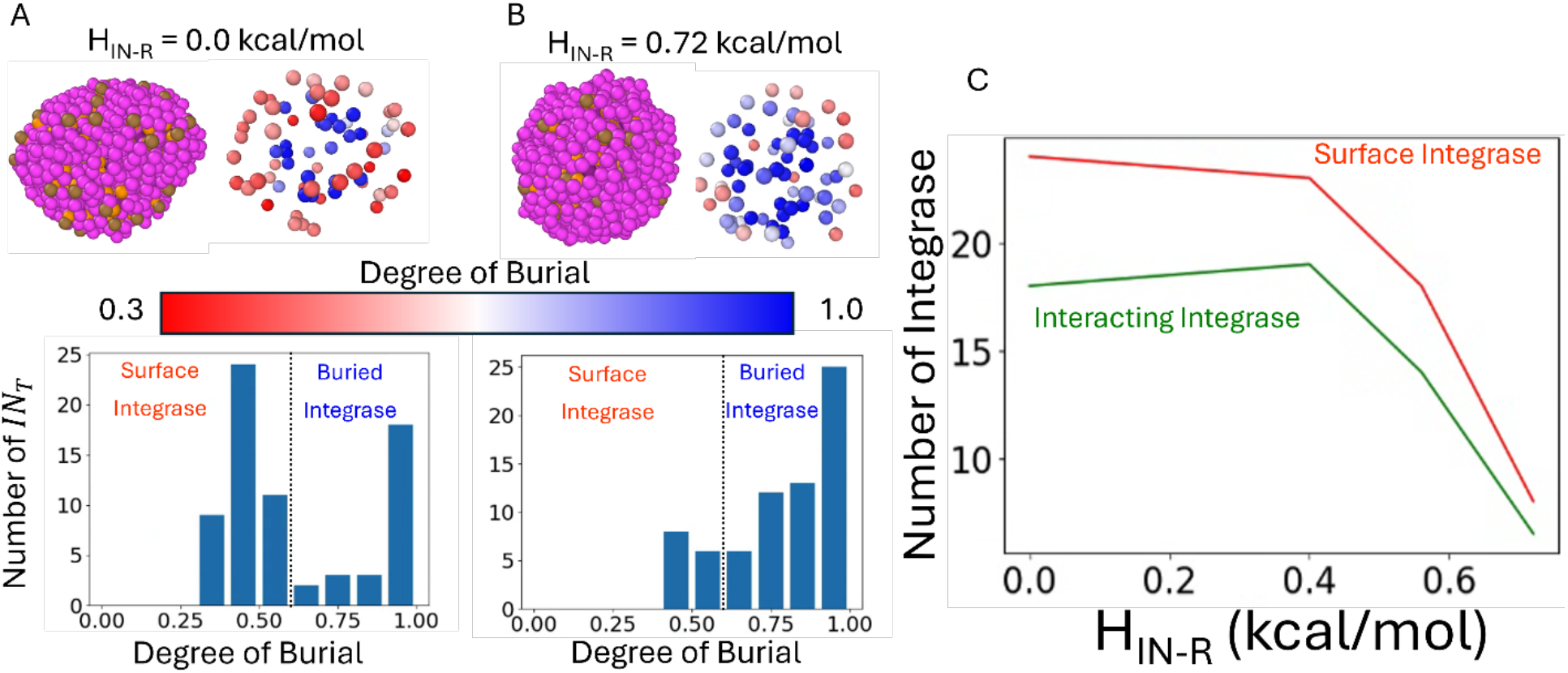
Partitioning of IN Interaction Sites between the Surface of the RNP and Buried within the RNP. Number of IN refers to the number of IN_T_sites (brown) which can interact with the CA proteins. There are 2 IN_T_sites per IN Tetramer and 35 IN tetramers in the RNP for a total of 70 IN_T_sites. (A) The RNP without attractive interaction between the IN and the rest of the RNP next to a visualization of just the IN_T_sites colored by their degree of burial. Below shows the bimodal distribution for all 70 IN_T_sites in this frame. (B) The same visualizations but with a strong attraction between the IN and the RNP. The bimodal distribution is shifted with far less surface IN. (C) The number of surface IN and number of INs that interact with the capsid proteins during assembly for a range of attractive interactions between the IN and RNP.

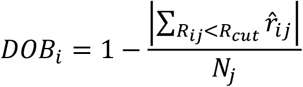

When the *IN*_*T*_bead was fully buried, its RNP neighbors were uniformly distributed, and the vector sum approached 0 and the DOB was 1. For a surface IN (*IN*_*T*_), the RNP neighbors were all on one side, and the vector sum was ∼ 0.5. Using this metric to evaluate all INs within the RNP complex, we observed a bimodal distribution of “buried” and “surface” IN. We chose a cut-off between the two categories to be 0.6, where a local minimum was observed.

This analysis allowed us to measure the number of surface INs as a function of H_IN-R_. In Figure 3C, we also saw the expected correlation between the number of surface INs and the number of *IN*_*T*_that interact with the capsid. This model allowed us to investigate how IN partitioning affects capsid assembly as opposed to the previous model where all the CA-RNP interactions were partitioned into a single domain. As shown in Figure 4, the stochastic capsid assembly process led to a distribution of assembled capsids even for the same value of H_IN-R_. Although they follow similar patterns of growth, random interjection of pentamers due to IP6 binding led to highly pleomorphic morphologies (see Figure S4). By comparing these distributions across different values of H_IN-R_, one can see how this parameter affects capsid growth. For the lowest value of H_IN-R_, we saw mostly spherical capsids (as defined in Figure S3) which are like the shapes observed in Figure 2B. Those capsids were observed for a uniform interaction between the capsid and the strongly condensed RNP. This makes sense, since the large amount of IN present at the surface also leads to a relatively uniform interaction across the RNP surface. In this case, we do not observe completely full capsids as in Figure 2, but the capsids tend to be smaller with more surface IN. In virus-based experiments, different sources quantify ensembles of capsids slightly differently. Ref. (67) classified 82% of capsids as cones and 18% as tubes, while ref. (28) classified 86% as conical, 12% as non-conical, and 2% as no core. Ref. (68) classifies ∼10% as immature, 25% as mature (tubular), 45% as mature (conical), and 10% as mature (irregular). Overall, this agrees best with the two cases with the fewest surface IN. Quantification of defects was done by ref.(68), with ∼25% of mature cores having a notable abnormality such as multiple cores present or multi-layering (onions). Smaller defects are likely unobservable in these studies. Ref. (23) observed the majority of their cores showing at least one defective local region, but this may have been related to purification procedures. They were able to quantify the numbers of hexamers in 6 complete cores (177, 182, 191, 198, 201, and 209) in reasonable agreement with our conditions with fewer surface IN.

**Figure 4.**
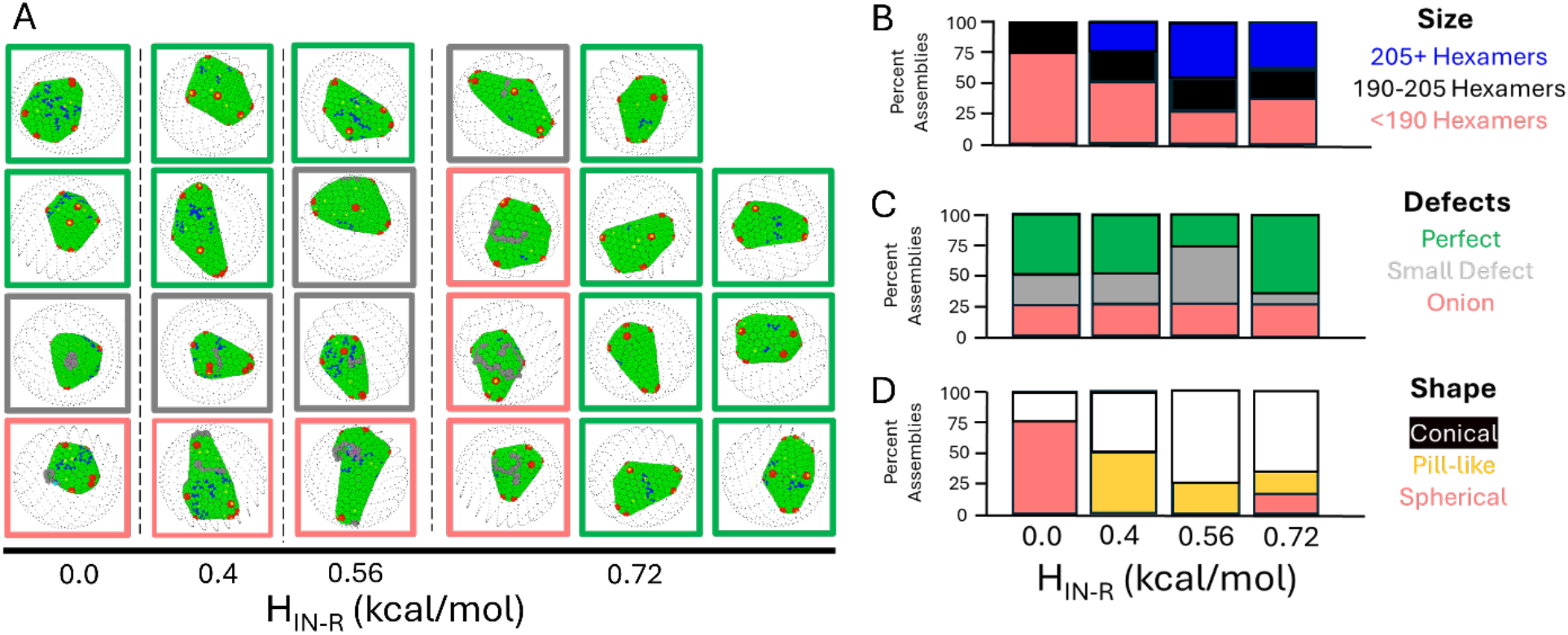
Gallery of Capsids. (A) Final assembly products shown for di=erent replicas of di=erent values of H_IN-R_. CA proteins shown in blue mean that they are interacting with IN_T_sites. One can see higher values of H_IN-R_ lead to fewer CA proteins interacting with the RNP through the IN_T_sites. (B,C, D) Bar charts quantify the distribution of di=erent assembly features (Size, Defects, Shape) as a function of H_IN-R_. The largest, most conical capsids with the fewest defects appear for high values of H_IN-R._

Since our RNP with IN tetramers appeared to be packageable, we wondered if it would be possible to for the presence of the capsid to induce filament assembly after or concurrently with the capsid assembly process. In adjusting our model of the RNP from the previously reported model(12), we did not remove the bonds tethering the IN tetramers to the RNP polymer, and thus they have limited mobility within the complex. However, we wanted to see if they would exhibit more complex organization without this restraint. Thus, we removed the bonds and some excluded volume within the RNP (see Supplementary Methods RNP Model 4 for details) to create an RNP with hypermobile IN that could respond to capsid assembly. We performed capsid assembly simulations in the same conditions and observed that the IN molecules arranged themselves in new ways in response to the capsid assembly. We observed that more IN partitioned to the inner surface as the capsid assembled and stayed there due to interactions with the assembling lattice (Figure 5C). Thus, in this case the RNP transitioned from having few IN tetramers to many IN tetramers on the surface during capsid assembly. This did not produce small or spherical capsids, as was the case when many IN molecules were present at the surface (see Figure 5B). We can explain this by noting that there is a lag time between capsid assembly and the recruitment of additional tetramers to the surface. Thus, IN was largely excluded from the surface, where it could have less impact the capsid assembly pathway. Instead, the capsid lattice growth impacted the behavior of the tetramers. We saw the tetramers organize themselves into bands or clusters on the RNP surface, rather than being randomly distributed as in the previous case (Figure 5A). As shown in Figure S5E, the average IN tetramer was in a cluster of 8 total tetramers. This seems to be due to the limited contact surface between the RNP globule and the capsid. The IN tetramers were only stable at the small section of the RNP surface to allow them to make hypothetical interactions with the capsid and thus the capsid controlled IN cluster formation. Despite this clustering long, straight IN-RNA filaments were not observed and using IN tetramers derived from the filament structures in Figure S5(C,D) did not reconcile this.

**Figure 5.**
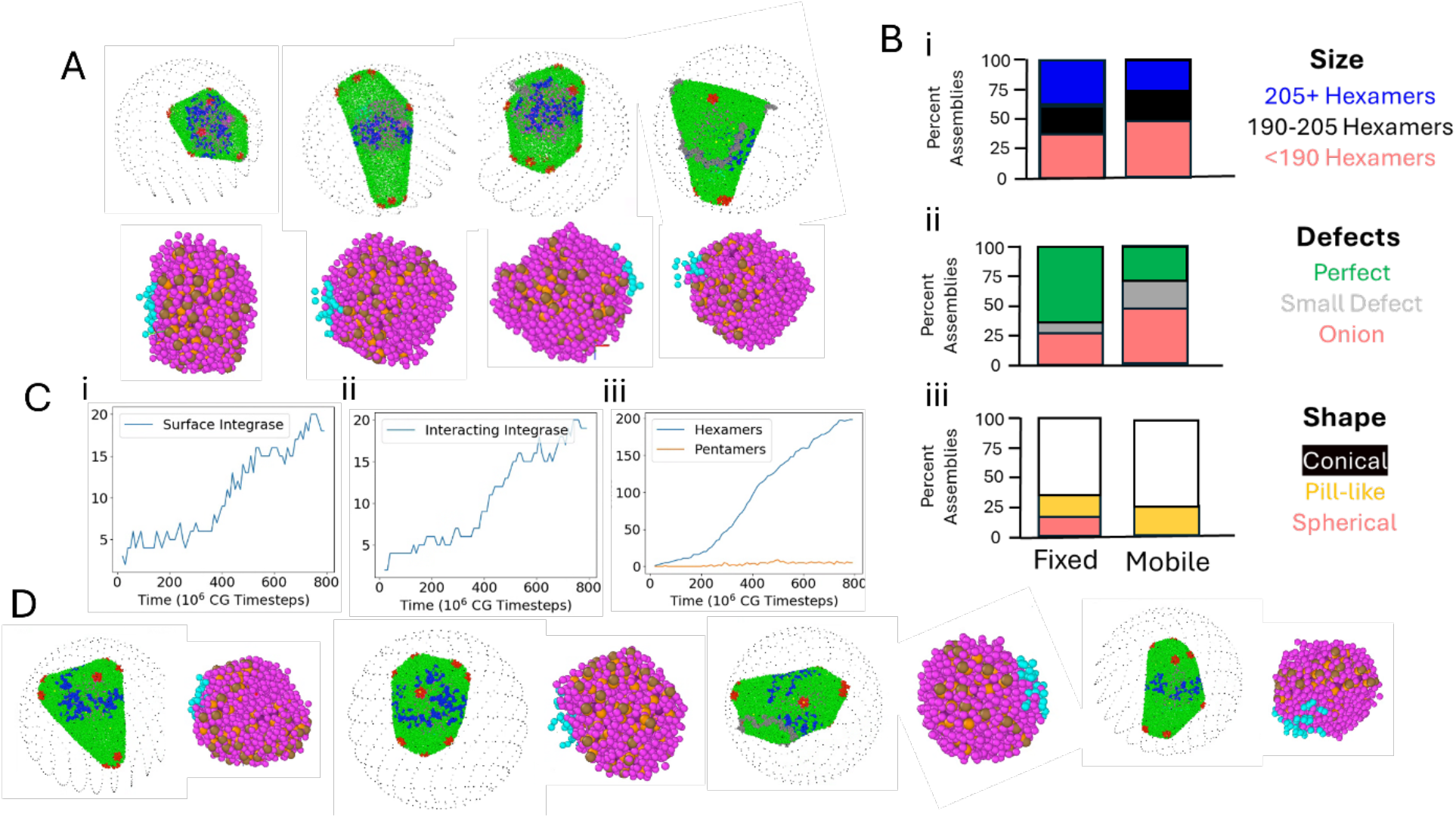
E1ects of Mobile IN on Capsid-RNA Assemblies. (A) Snapshots of end-point capsids resulting from simulations where the INs were hypermobile within the RNP are shown on top of snapshots of the RNP itself, which are enlarged to view the IN_T_sites more visible. The RNPs show many IN_T_sites on the surface clustered together. (B) Bar charts compare the di=erent assembly features between the H_IN-R_=0.72 kcal/mol case and hypermobile IN case shown in (A). The distributions are relatively similar. (C) The time evolution of the surface IN, interacting IN and hexamers/pentamers show that IN arrives at and stays at the surface of the RNP due to interactions with the capsid. Thus, IN assembly at the surface lags capsid assembly. (D) Snapshots of capsids and corresponding RNPs when we imparted IN mobility after capsid assembly was completed. The same trend as in (A) was observed with clusters or streaks of IN appearing at the surface of the capsid. The finished capsids come from Figure 4(A).

We also wanted to test the case where IN-RNA filament assembly would occur completely after the capsid was assembled. To test this, we ran a set of simulations where we took four final configurations from the low surface IN cases and switched the model to allow the IN to become hypermobile. We then relaxed these configurations and ended up with the final structures shown in Figure 5D. They have similar clustering characteristics (the average tetramer is in a cluster of over 5 tetramers) as the cases where IN tetramers were mobile during assembly in Figure 5A and thus it seems that IN mobility does not have a large impact on the final structure of these clusters. We also tried to create a model where IN-RNA filaments could form because of the capsid assembly growth on the RNP. Details on this model can be found in the Supplementary Information for Figure S5 and RNP Model 5. However, we did not see clear evidence that the capsid can induce self-assembly of long IN-RNA filaments that would not assemble without the influence of the capsid.

### Octameric IN-RNA filaments template the formation of biologically relevant HIV-1 capsids

We then decided to explore how the capsid assembly would be affected if *we assumed that octameric IN-RNA filaments form prior to the initiation of assembly*. We developed a CG model of these filaments which is described in more detail in the Supplementary Methods RNP Model and Figure S7-S10. The model was built by first running all-atom molecular dynamics simulations of an IN 16-mer, composed of two octamers, using a recently published experimental structure(45). We next built a hetero-elastic network (HENM) model(69) of the 16-mer using intra and inter-octamer bonds, which allowed us to have an HENM for a filament of arbitrary length. We then connected this model to the same RNP polymer we used in the studies of IN tetramers and employed angle potentials on the polymer to enforce the rigidity of the filaments, which has been experimentally reported(45). We used a relaxation process to produce our final RNP model to initiate assembly. Since we could create filaments of any length and add any number of filaments to the RNP, we first sought to explore how the number and length of filaments impacted assembly. We had previously assumed 35 IN tetramers were present in the RNP, which is 140 total units of IN, consistent with experimental measurements(21). In Figure 6A and B, we investigated cases of 5 filaments, which are each 3 octamers long and 3 filaments, which are each 5 octamers long. This gives a total of 15 octamers or 30 IN tetramers in each case, which is like our previous assumption. When we assembled the capsid around these RNPs, we found that they were comparatively small (see Figure 6A and B), consistent with observations from the many IN tetramer case(H_IN-R_=0.0). Considering that these RNPs contained even more surface IN tetramers (30), this reinforced the idea that high levels of surface IN led to smaller capsids. One difference was that the shape of these capsids was more conical and less spherical than we had observed with just surface IN tetramers (Figure 4D, H_in-R_=0.0). *This first suggested that the rigid IN-RNA filaments could help to template conical capsid structures*.

**Figure 6.**
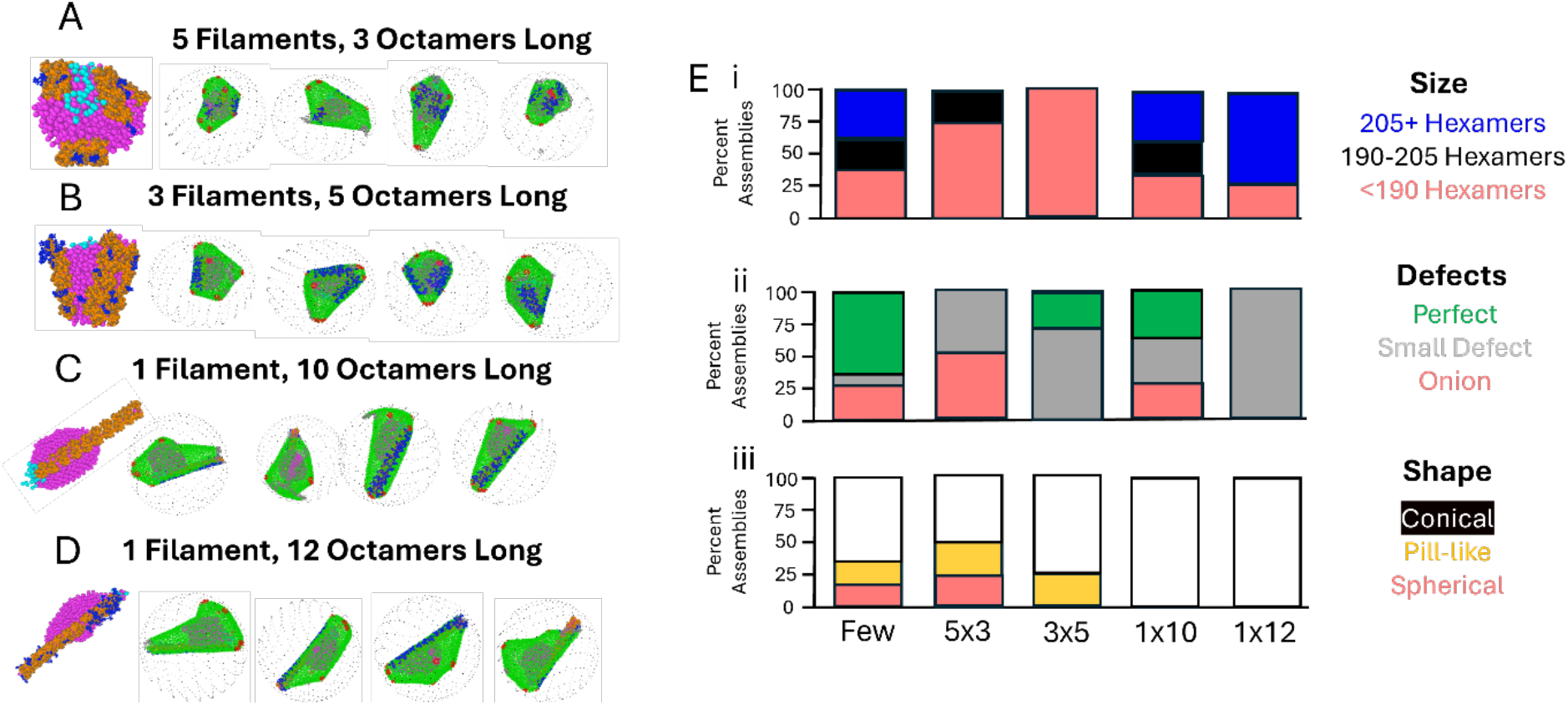
Capsid Assembly with octameric IN-RNA filaments. (A) Snapshots of capsids assembled around the RNP with 5 filaments, 3 octamers long shown in the upper left. (B) Snapshots of capsids assembled around the RNP with 5 filaments, 3 octamers long shown in the upper left. (C) Snapshots of capsids assembled around the RNP with 1 filament, 10 octamers long shown in the upper left. (D) Snapshots of capsids assembled around the RNP with 1 filament, 12 octamers long shown in the upper left. (E) Comparison of the size, shape, and defects of the capsids assembled in cases (A-D) as well as Figure 4 with limited surface IN tetramers.

We decided to consider the case of only one longer filament based on our earlier observations with IN tetramers and also from the experimentally-derived filament structure (40). We introduced filaments that were 10 and 12 octamers in length so that they would not be too long to fit in the capsid/virion but would also take up most of the available 35 IN tetramers. The 10 and 12 octamer filaments are composed of 20 and 24 tetramers, respectively. When we assembled the capsid in the presence of these single filament templates, we observed much larger capsids that are more in line with experimentally-determined capsid sizes than the cases initiated with multiple filaments. We did notice that some of the capsids had “busted tips”, characterized by a small gap at the end and less than 12 pentamers. These have been observed experimentally (70), but we observed that this was the behavior especially for the filaments of length 12. It appeared that such long filaments combined with the virion envelope boundary made it difficult for the capsid to have enough room to close. Interestingly, there is often space for a longer filament in closed capsids with a filament of length 10 in Figure 7D and E. This lack of room to close may also be related to the present modeling of the virion as a rigid sphere (see Supplementary Methods for more details), while the real virion membrane is likely more compliant and aspherical. Overall, one filament of length 10 matches the previously discussed experimental capsid ensembles the best, with the caveat that this model is not producing tubes/pill-like capsids. It does produce elongated conical morphologies, but our RNP globule appears to be too tightly condensed to fit in a narrow tube capsid.

**Figure 7.**
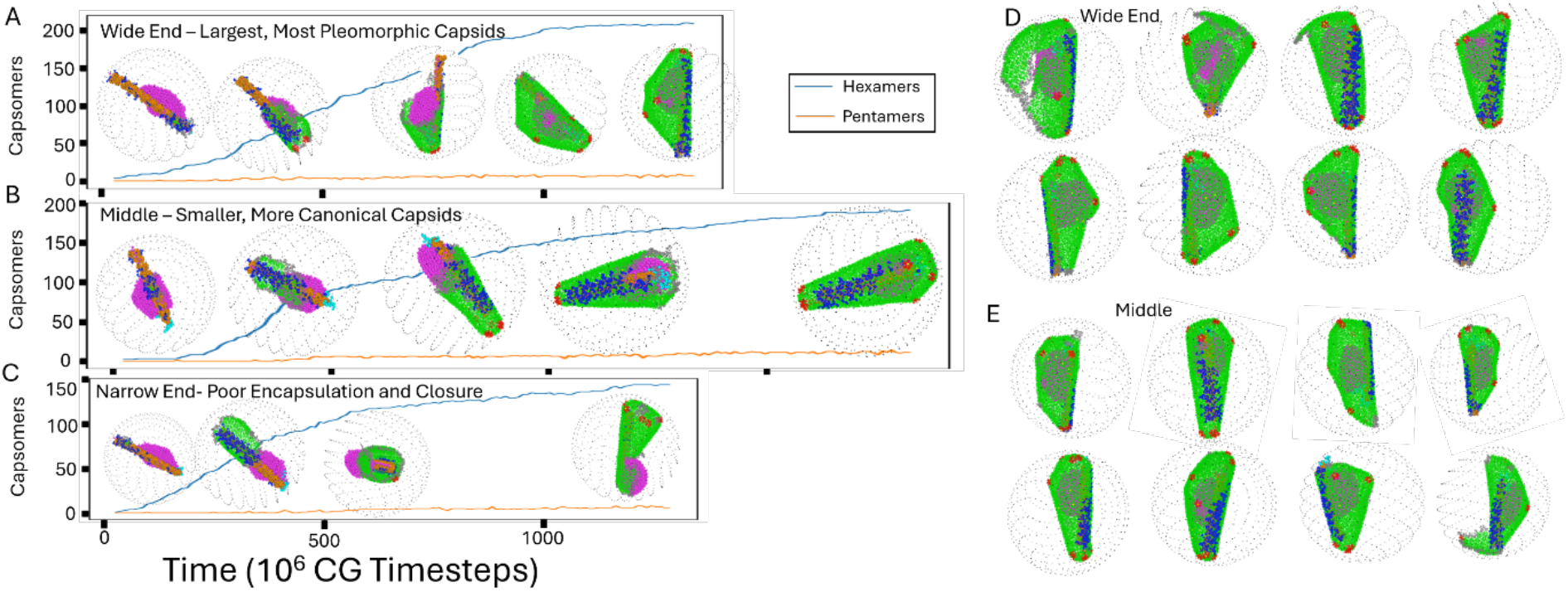
Capsid Assembly Kinetics with Octameric IN-RNA Filaments and DiDerent Initial Growth Sites. (A) One representative capsid assembly pathway when the initial filament binding occurred at the wide end of the RNP. The capsid lattice quickly grew around and encapsulated the RNP globule and then proceeded down the filament, resulting in a large capsid. (B) One representative capsid assembly pathway when the initial filament binding occurred at the middle of the filament. The capsid lattice quickly grows down the filament, encapsulating the narrow end. Growth at the wide end was slower and the resulting capsid was conical. (C) One representative capsid assembly pathway when the initial filament binding occurred at the narrow end of the RNP. A tubular structure formed at the narrow end without encapsulation of the globule, leading to stalled growth. (D) Gallery of capsids for growth nucleating at the wide end of the RNP. (E) Gallery of capsids for growth nucleating at the middle of the filament.

When modeling the RNPs with one large filament we also assumed that the filament should initiate at the 5’ UTR since there is experimental evidence that the filaments prefer TAR RNA found in the 5’ UTR. In the case of many filaments, we placed them randomly, since they would all have to start growing from different locations. Under this assumption, the RNP became highly asymmetric with most of the excess length of the filament in one direction from the globule, as shown in Figure 6C and D. If the filament initiated outside of the 5’ UTR, its ends were symmetric (Figure S10). Thus, the RNP develops a “wide end” and a “narrow end”, akin to the conical capsid. While assembling the capsids around this RNP, we noticed that we would get different results based on where the initial hexamer bound on the filament. We classified the binding site locations as the wide end, middle, and narrow end.

For all three growth initiation sites shown in Figure 7, we observed that the capsid lattice started to grow along the filament and avoided the largest parts of the globule. For the middle and wide end, the capsid must grow around the globule, but for the narrow end, a tube-like structure often formed before it “reached” the globule and thus growth exterior to the globule finished quickly while growth around the globule was slow or completely stalled. (see Figure 7C). Evidence of this disjointed growth may be visible in Figure 8(B). Although it is not possible with our seeded capsid growth approach, this stalled growth may be an opportunity to nucleate and grow a second capsid, which has been observed experimentally(68). For the wide end case, encapsulation was comparatively simple and occurred before a tube formed at the narrow end. This started the capsid on a path where the wide end could experience greater pleomorphism (see Figure 7A, D). For the middle case, we saw the narrow end close first, but the RNP was still encapsulated. From here it slowly closed around the globule, resulting in the most conical capsids among all our simulations, which were smaller than the wide end capsids (see Figure 7B, E). This follows a previously proposed model wherein the capsid lattice “grows rapidly along one side toward the narrow tip, followed by slower growth of the structure around the body of the cone”(16). *Our simulations suggest that the IN-RNA filament directs this asymmetric growth*.

**Figure 8.**
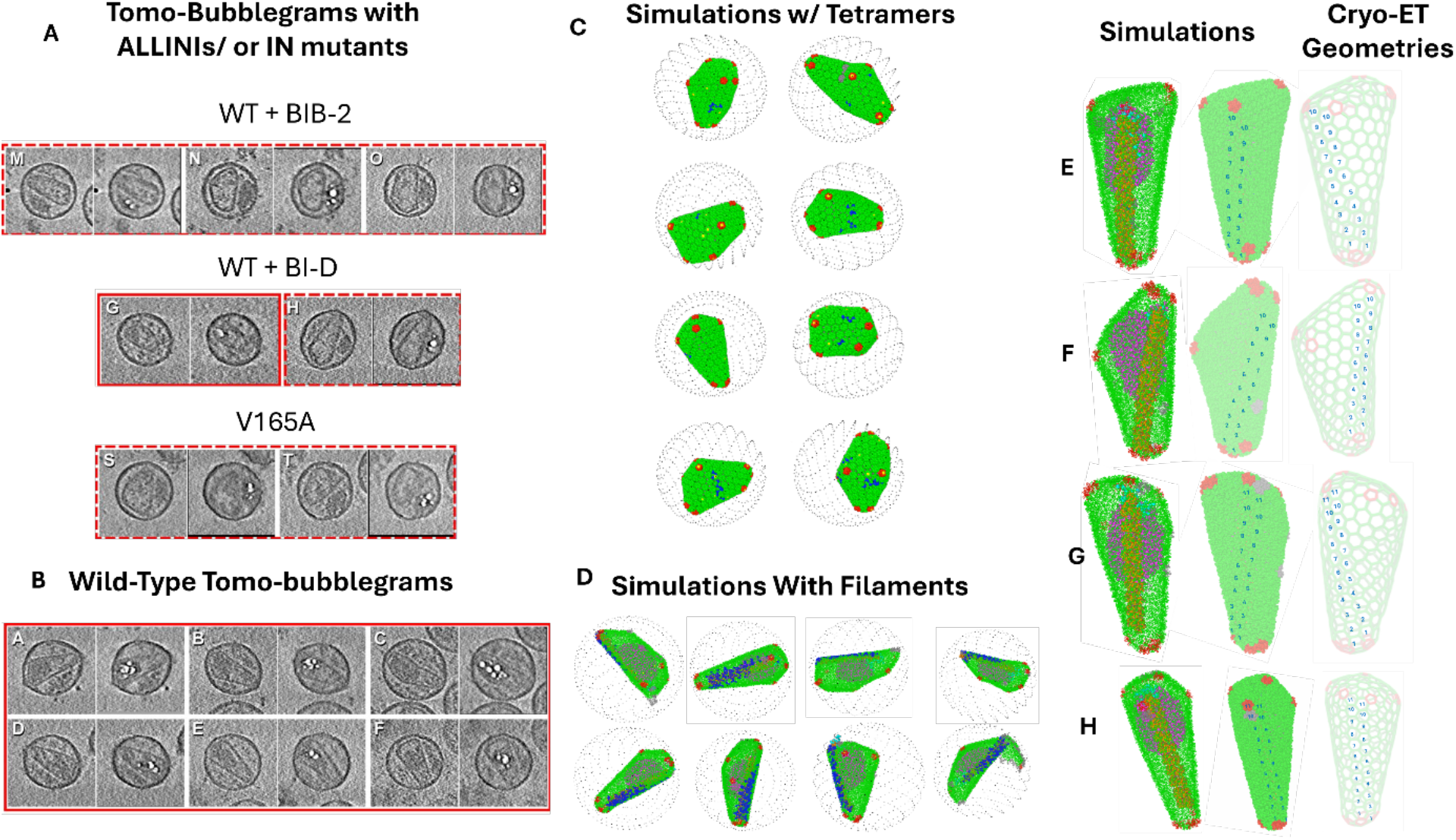
Comparisons across Experiments. (A, B) Tomo-bubblegrams of HIV-1 cores classified as conical in the original manuscript(28) (adapted from Fontana et al., J. Virol. 2015 with permission from the American Society of Microbiology). BIB-2 and BI-D are two different ALLINIs; V165A is a class II IN mutant virus. (C) Assembled capsids with the tetramer RNP are less elongated as capsids assembled in the presence of IN-RNA filaments, shown in (D), just as cores with disrupted IN-RNA binding in (A) are less elongated as the wild-type cores shows in (B). (E-H) Comparisons of our assembled capsids to solved lattices from previous work(23, 24). The first structure shows our assembled capsid with the IN-RNA filament and RNP visible. The second shows the long runs of flat hexamers that form because of the filament, and the last shows a similar experimental morphology with the run of flat hexamers numbered in the same way.

As shown in Figure 8, the capsids assembled alongside IN-RNA filaments (Figure 8D) are noticeably elongated compared to those that assembled around the RNP with IN tetramers(Figure 8C). This difference bears a strong resemblance to tomo-bubblegrams from previous work, showing that wild-type viral cores (Figure 8B) are far more elongated than those where IN-RNA binding was disrupted through ALLINI treatment or the class II IN mutation (Figure 8A). This suggests that the IN-RNA filament may help to elongate the HIV-1 cores. Additionally, the capsids assembling on the filaments had long regions of at least 10 flat consecutive hexamer pairs due to the rigidity of the filament. In Figure 8(E-H), we number these long runs of flat hexamers on our assembled capsids and show they also exist in experimental cryo-ET structures(23), which bear a striking resemblance to our assembled capsids. These long flat runs were observed in only approximately 10% of the structures when we assembled around the RNP with IN tetramers (Figure 8C). Moreover, five of the six complete capsid structures from a previous publication(23) showed these long runs. Outside of HIV-1, internal scaffolding is often used to guide capsid morphology(71) and elongation of protein shells by internal scaffolding has been observed in bacteriophage(72).

## Discussion and Conclusions

In this study, we investigated several models for how the RNP could impact HIV-1 capsid assembly. In some ways, these models showed similar behavior. Regardless of how we modeled the RNP component to interact with the capsid, the organization of IN within the RNP controlled the resulting capsid assemblies. When there was too much IN scattered across different regions of the RNP globule surface, this led to smaller capsids with less free interior volume for reverse transcription. This behavior was especially obvious in the simplest RNP model, where we modelled IN implicitly. When we included explicit IN tetramers, we observed that having only a few surface tetramers at the beginning of assembly was important to assemble a capsid that was large enough to accommodate DNA synthesis. However, it was possible to have many tetramers at the end of the assembly, since our hypothetical interaction between CA and the tetramer could hold mobile tetramers at the RNP surface. When the tetramers reached the RNP surface, they accumulated within a small region touching the capsid, but this clustering did not lead to long filament structures or capsids with the long runs of hexamers that would interact with them. We further investigated capsid assembly in the presence of preformed octameric IN-RNA filaments and found that a single long filament produced much more elongated morphologies featuring these runs of 10 flat hexamers. This is consistent with previous experimental images(28), which showed that ALLINI-treated or class II V165A mutant virions, where IN-RNA binding and IN oligomerization is disrupted, were much less likely to form elongated conical capsids.

Since the kinetics of assembly processes play such an important role, it will be important in the future to better understand the different kinetic aspects of maturation. This includes Gag (and Gag-Pol) processing, which precedes assembly and encapsulation, and the two should be closely linked. Slow cleavage of CA relative to IN would give more of an opportunity for filament assembly to progress before capsid assembly, while faster cleavage would give less time for filament assembly before capsid assembly starts. The cleavage rate of NC would also be very important in condensing the RNA, as well as any changes to the base CA assembly rate(13). Modeling the entirety of this process can increase our understanding and help to inform experiments given the difficulty of imaging and resolving in time these processes.

The next steps in the HIV-1 replication cycle, reverse transcription and uncoating, are also kinetic processes that will be impacted by the end structures we observed here. For instance, a highly structured RNP containing an accessible 5′ UTR, as we suggest, could play a crucial role in organizing reverse transcriptase to initiate reverse transcription. It has also been noted that the filament structures of length 2 are very similar to intasomes, which must form for DNA integration(51); but how the filaments may break apart into these smaller structures is currently unknown and may be linked to rupture of the capsid lattice. Alternatively, the filament could stabilize the capsid lattice and redirect rupture to specific sites by preventing dsDNA from pushing on the inner capsid wall(30) or mechanically stabilizing it. Further simulation work could help to expand upon these hypotheses.

In conclusion, we investigated two different hypotheses for the organization of IN within the RNP. Both could lead to cores with sufficient volume for reverse transcription but with different characteristics. Assembly with IN tetramers did not lead to the formation of any long IN-RNA filaments and that these capsids were wider despite having nearly the same number of hexamers. In the case of the preformed IN-RNA filaments, we saw that the most experimentally consistent structure was one long filament and that the resultant capsids were highly consistent with elongated conical capsid morphologies featuring runs of roughly 10 flat hexamers observed in experiment. Thus, we suggest the IN-RNA filament forms ahead of the capsid and directs it to its conical morphology. As there is growing evidence that the wider capsids cannot squeeze through the nuclear pore efficiently and enter the nucleus for integration(4, 67), this suggests the IN filament may be important for infectivity.

## Methods

### CGMD Simulations

Details on the CG models for CA, RNP models including IN, IP6, and the crowding agents are provided in the Supplementary Information. Self-assembly of HIV-1 CA was simulated using CGMD in the canonical (constant NVT) ensemble at 300 K with Langevin thermostatting with a 10fs timestep and a damping period of 100ps. In the setup for Figure 2 without explicit IN, an 88 nm × 88 nm × 88 nm cubic box with periodic boundaries and with 750 CA dimers, 1700 inert crowders, 521 IP6, and one RNP is prepared using (https://github.com/waltmann1/CGBuilder). In the virion-like environment used in all simulations with explicit IN, a virion-like boundary condition is created by placing particles on the surface of a 144 nm sphere to match the approximate size of the virion. The area per particle is 12 nm^2^ /particle and they interact uniformly with all other species such that all particles are confined to the sphere. This doubles the volume and as such all species are doubled except for the RNP. The increased number of capsid monomers increases assembly rates and thus the fraction of active CA monomers ([CA]_+_/[CA]) decreased from 0.15 to 0.10 to observe proper assembly. This value is more in line with experimental observations in vitro(22). For efficient sampling of nucleation events, simulations were initialized with a much higher fraction of active CA and then trajectories yielding a nucleated hexamer were selected to seed production CGMD simulations.

### Data Analysis and Visualization

Graph analysis of assembled lattices was performed using the Python package NetworkX 2.1 (http://networkx.github.io/). We create a graph of CA monomers with two different types of edges, N-terminal domain edges and C-terminal domains. NTD edges are defined by the NTD dimer interface, i.e., when the distance between CG types 132/31 or 133/33 was less than 2.0 nm. CTD edges are defined between each pair of monomers in a dimer. By considering only the NTD edges we get small subgraphs that represent the oligomers present in the lattice. When these subgraphs have 5 nodes and those nodes have 10 neighbors total, that is considered a pentamer. Most hexamers are subgraphs with 6 nodes and 12 neighbors. However, we observe some hexamers where a free CA monomer has made one loose contact with a monomer in the hexamer. Thus, some hexamers have 7 nodes and 5 of those nodes have 2 neighbors, 1 node has 3 neighbors, and 1 node has only 1 neighbor. All other subgraphs are defined as disordered CA.

## Acknowledgements

Research reported in this publication was supported by the Behavior of HIV in Viral Environments (B-HIVE) Center of the National Institute of Allergy and Infectious Diseases (NIAID) of the National Institutes of Health (NIH), under award numbers U54AI170855 (50%), R01AI178850 (50%) to G.A.V., U54AI170855 (A.N.E.), and R37AI039394 (A.N.E.). C.W. acknowledges the support of an NIH Ruth L. Kirschstein National Research Service Award Postdoctoral Fellowship under award number F32AI186429. The content is solely the responsibility of the authors and does not necessarily represent the official views of the NIH. Computational resources were provided by the University of Chicago Research Computing Center and the NIH-funded Beagle-3 computer (NIH award 1S10OD028655-01). We would also like to acknowledge Peter Cherepanov and Florian Chenavier for helpful discussions, generously sharing experimental data, and comments on the manuscript.

## Author Contributions

C.W. M.G. and G.A.V. designed research; C.W. performed research; C.W., M.G., A.E., D.L. and G.A.V. analyzed data; C.W. methodology; G.A.V. project administration, funding acquisition, resources, supervision; and C.W., M.G., A.E., D.L., and G.A.V. wrote the paper.

