## Supplemental Methods and Figures 1-11 for "HIV-1 Ribonucleoprotein Complex Organization Facilitates Assembly of the Mature HIV-1 Capsid"

### Supplementary Results

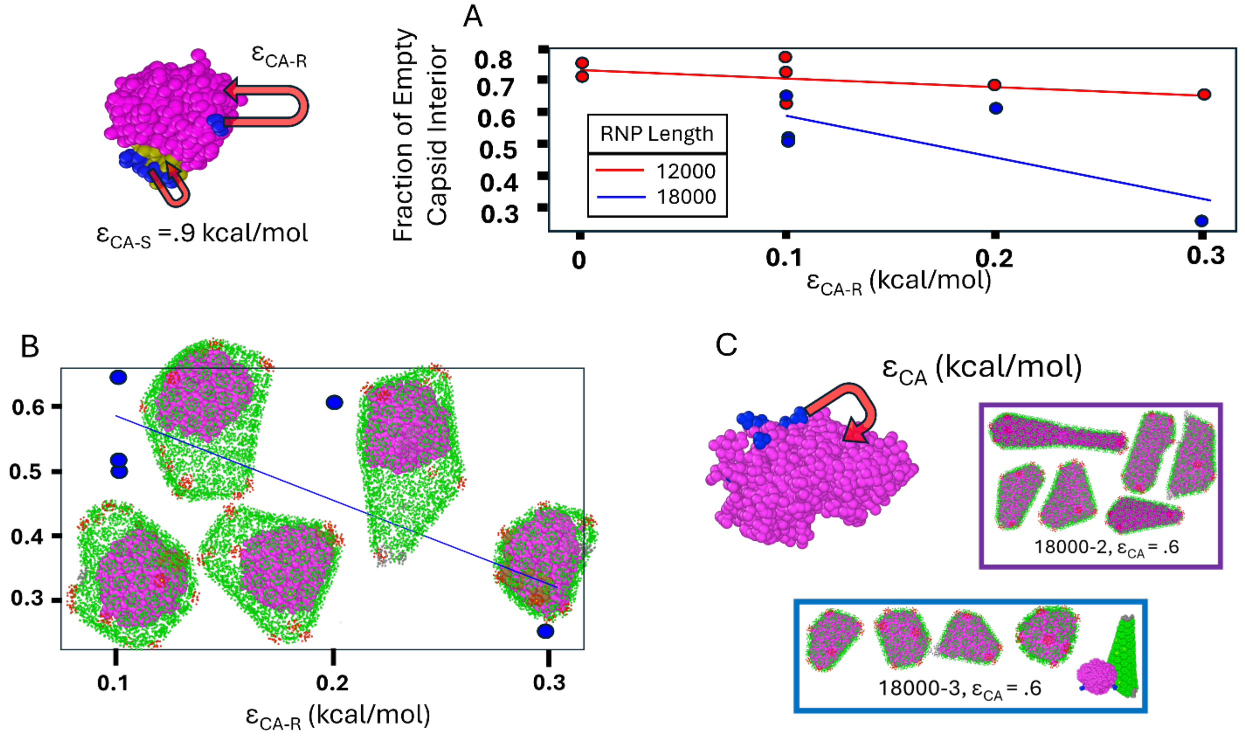


***Figure S1.*** **One-component RNP model details.** (A) The fraction of empty capsid interior is shown as a function of $\varepsilon_{CA-R}$, the interaction between capsid and the rest of the RNP, for 12000 and 18000 CG bead long RNPs. The CA interaction with the packaging domain or signal is a constant 0.9 kcal/mol (left). (B) Snapshots of the capsid-RNP complex are shown for the 18000 length RNP visualized such that the RNP and empty space in the capsid interior is visible. (C) More replicates are shown for single component RNPs with parameters where capsids form completely full. These RNP-containing capsids are completely full, and this is inconsistent with experimental results as the RNP could not complete reverse transcription within these capsids.

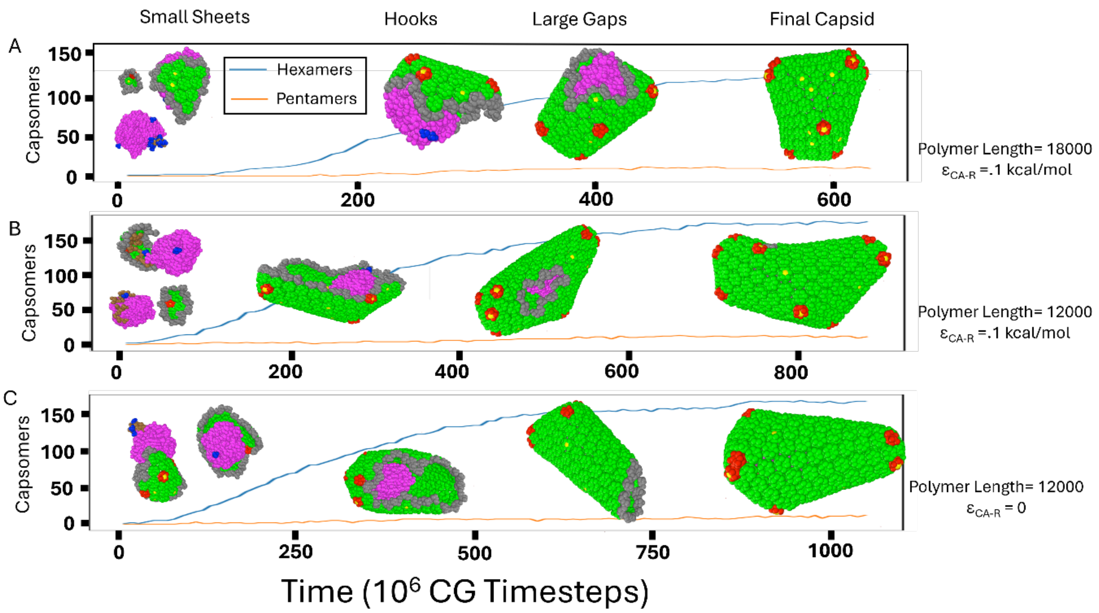


***Figure S2.* Assembly pathway of capsid assembly around the RNP.** Trajectories of capsid assembly are shown by time series data of the number of hexamers and pentamers present in the growing CA lattice as well as snapshots. These snapshots show the 4 major intermediate states in capsid assembly: small sheets, hook-like structures, capsids with large gaps, and the completed capsid. Parameters of the RNP are shown to the right for (A) polymer length 18000, $\varepsilon_{CA-R}$=0.1 kcal/mol (B) polymer length 12000, $\varepsilon_{CA-R}$=0.1 kcal/mol, and (c) polymer length 12000, $\varepsilon_{CA-R}$=0 kcal/mol.

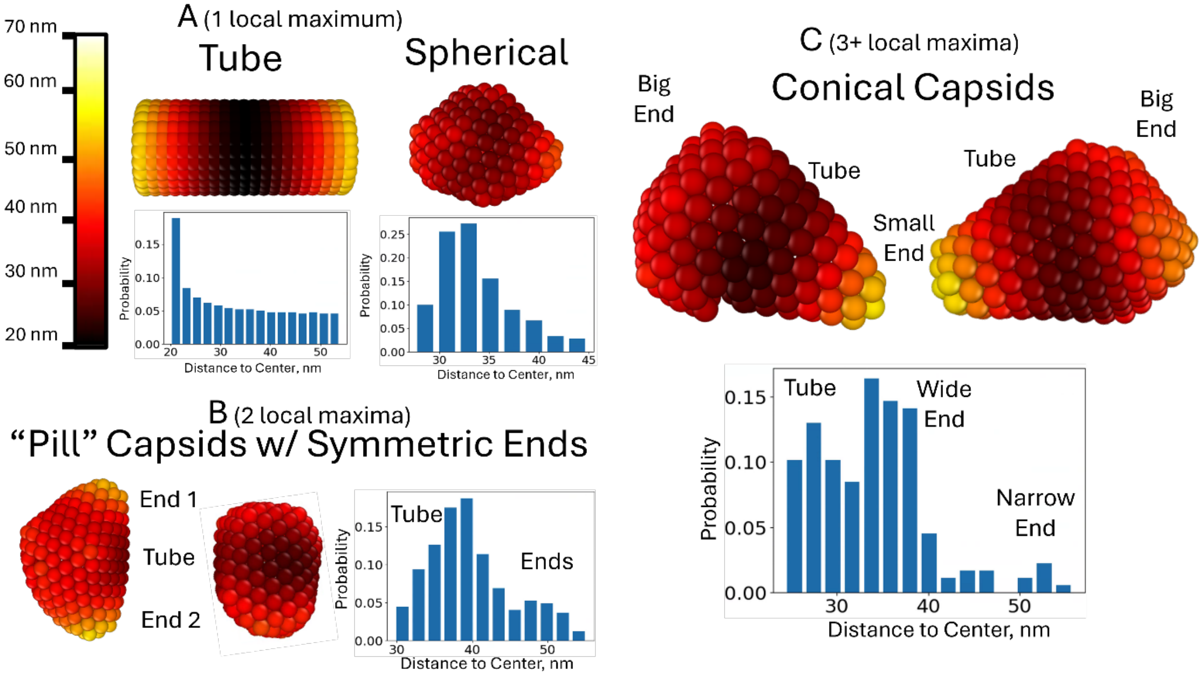


***Figure S3. Capsid shape is quantified by the distribution of distances between the centers of CA oligomers (including pentamers and hexamers) and the geometric center of the capsid (the average position of all CA oligomers).*** *(A,B) When this distribution has a single maximum, the shape tends to be either a sphere or tube. Individual beads show the positions of CA oligomers and are color coded by distance, such that they can be related to the full probability distribution shown below. (C) Examples of “pill” capsids that have two maxima related to the tube and symmetric ends of the capsid. (D) Examples of conical capsids featuring 3+ local maximums related to the tube and the larger and smaller ends of the capsid.*

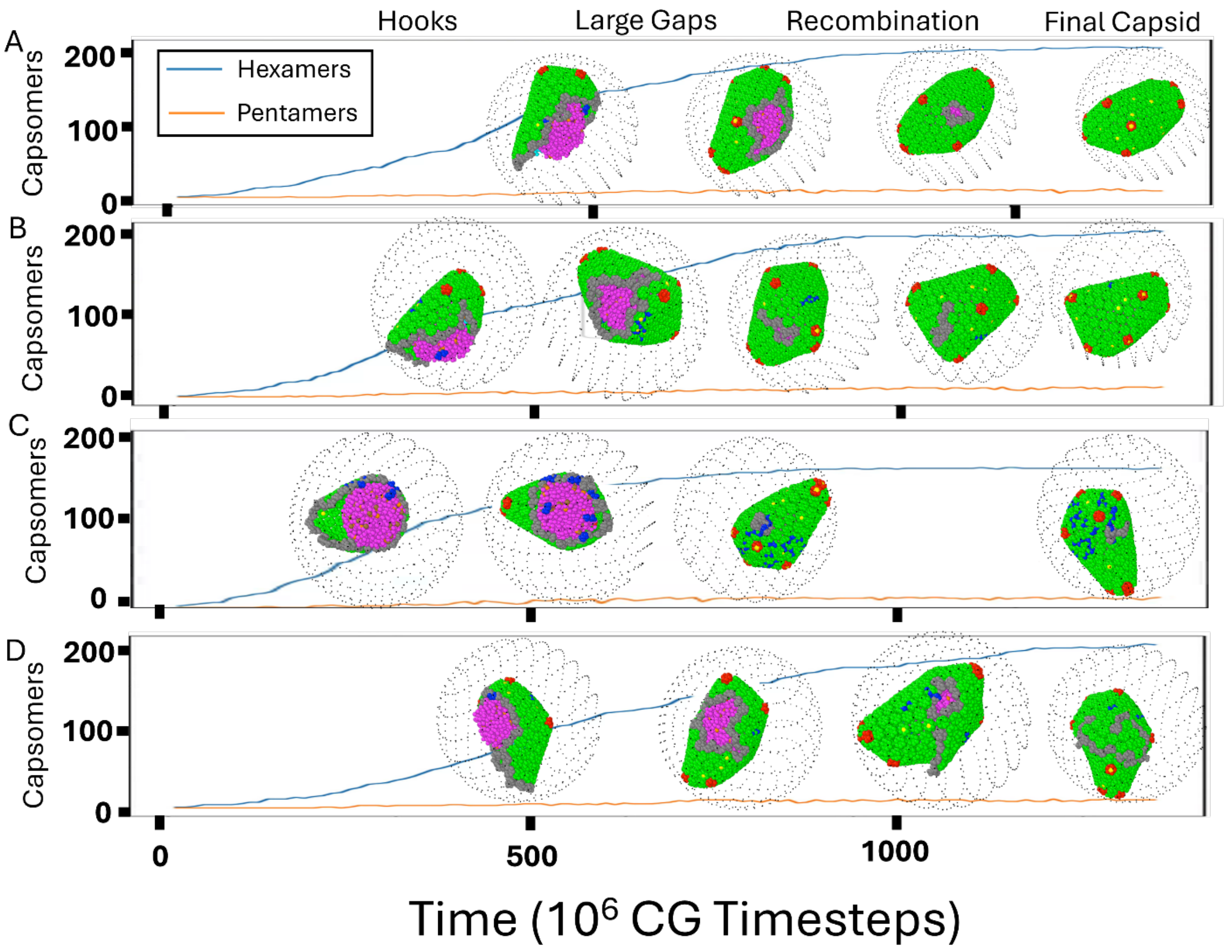


***Figure S4. Assembly Kinetics and Closure.*** *Time-series data for the addition of hexamers and pentamers are shown for four representative cases of assembly along with simulation snapshots displaying Hooks, Large Gaps, Recombination and the Final Capsid. (A,B,D) show cases where* $H_{IN\text{-}R}$ *= 0.72 kcal/mol while (C) shows* $H_{IN\text{-}R}$ *= 0.56 kcal/mol. (A,B) show cases of successful closure of a capsid although the process is slower in B due to a slow to heal seam in the capsid lattice. (C) Shows a small defect in the capsid related to interactions with between the capsid proteins and surface integrase. There are more surface integrase present in this case since* $H_{IN\text{-}R}$ *is lower. (D) Shows a case of multilayering that occurs in during failed recombination.*

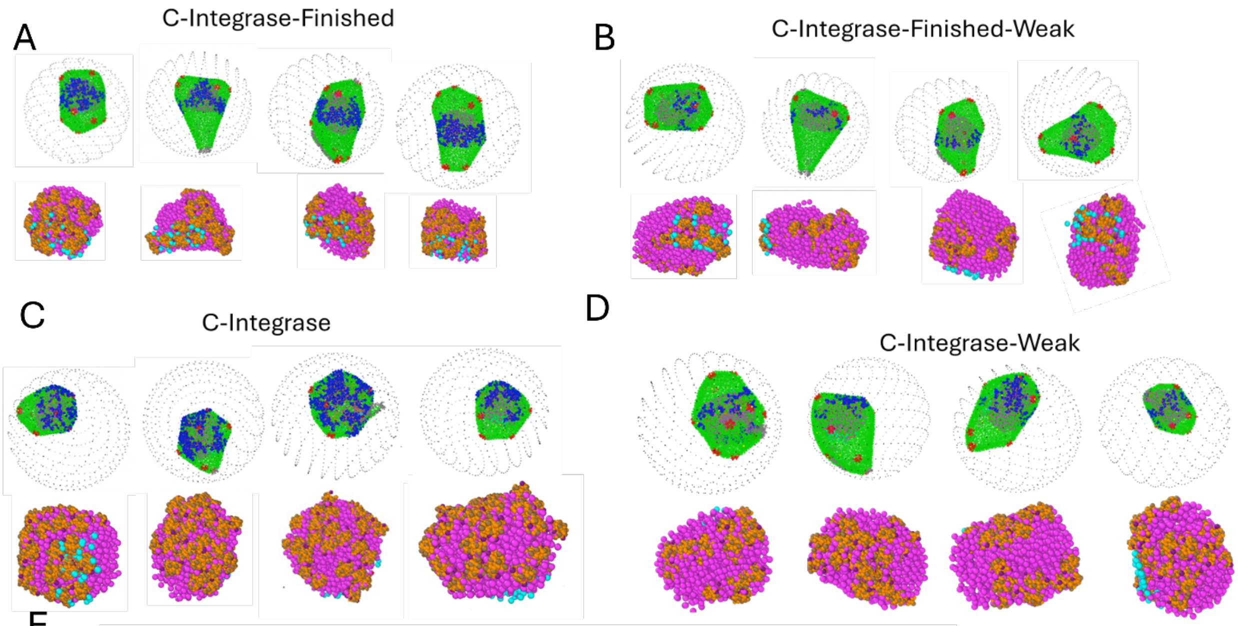


E

| | L | L-Fin | C | C-Fin | C-Weak | C-Weak-Fin |
|---|---|---|---|---|---|---|
| Surface IN | 19 | 19 | 35 | 34 | 22 | 21 |
| Interacting IN | 19 | 19 | 35 | 34 | 20 | 17 |
| Cluster MW | 8.0 | 5.2 | 5.5 | 7.1 | 2.5 | 2.2 |
| Filament MW | N/A | N/A | 2.7 | 2.9 | 1.6 | 1.5 |

***Figure S5. Assembly results with C-Integrase capable of forming octameric filaments through tetramer-tetramer interactions.*** *These tetramer-tetramer interactions are chosen such that filaments will not form in the absence of capsid. The goal is to test whether the capsid can induce filament formation. The case of filaments that form independent of the capsid is tested extensively in the main text. (A) Starting with a finished capsid C-Integrase form clusters at the surface of the RNP. Blue shows the part of the capsid interacting with the integrase and the figures below show just the RNP without the capsid. (B) The interaction between CA and the C-Integrase is weakened leading to less C-Integrase at the surface and thus less clustering although the shape is similar. (C) Capsids are assembled with the highly mobile C-Integrase in the RNP leading to spherical capsids due to C-Integrase affecting the ability of capsid to grow off the RNP as has been seen previously with the uniform RNP models in Figure 2. (D) The interaction between CA and C-Integrase is weakened and thus the capsid growth is not as affected by C-Integrase. There is less C-Integrase on the surface and less clusters (E) Table summarizing the number of IN on the surface of the RNP and Interacting with the capsid out of 35 total. L-IN is the tetramers in the main text. The Cluster MW is the weight average molecular weight of the clusters which can be thought of the size of the cluster that the average C-Integrase tetramer is in. The Filament MW is the same as the Cluster MW except only interactions between the attractive sites count towards determining what is a cluster rather than proximity. See Supplementary Methods for more details. Overall, these results do not support the idea that filaments only form in response to templating from the capsid. They are not opposed to the idea that the capsid can encourage growth of filaments.*

**Supplementary Methods**

<u>*Coarse-Grained Model Details*</u>

**Capsid Protein (CA)**

In this study, we used the coarse-grained model of the HIV-1 CA dimer that was previously developed in our group(*1, 2*). Here, we will note some of the key features of the model although the full description is available in those works(*1-4*). The model is based on 2 separate elastic network models of the N-terminal domain and C-terminal domain of the CA monomer. These two elastic networks use the harmonic potentials in Eq. S1 and K=1000 kcal/mol/nm$^2$.

$$U_{Bond}(r) = K\,(r - r_0)^2 \qquad S1$$

The two domains are connected by a flexible linker domain where K=1 kcal/mol/nm$^2$. Interactions which hold together CA monomers and allow them to assemble are given by double gaussians potential shown below.

$$U_{Bind}(r) = -A\exp(-Br^2) - C\exp(-Dr^2)]\,, r < r_{cut} \qquad S2$$

The values of the four parameters are A = 1.1 kcal/mol, B = 10 nm$^{-2}$, C = 2.0 kcal/mol, and D = 100 nm$^{-2}$. The potential is applied for specific “ghost” sites which are embedded in the CA monomers but have no excluded volume interactions and do not contribute to the shape of the monomer. These interactions can be enabled and disabled to create the $[CA]_+$ and $[CA]_-$ populations.

***RNP Model 1 (1 Component)***

RNP Model 1 is the simplest model of the RNP. It is a bead spring model of a linear polymer with different lengths (6000,12000, and 18000 CG beads). A harmonic potential is applied across consecutive monomers.

Where K, the harmonic spring constant is 1000 kcal/mol/nm$^2$. And the equilibrium distance, $r_0$, is 1.25 nm. Interactions between polymer beads are given by a Lennard-Jones potential:

$$U_{LJ}(r) = 4\varepsilon\left(\left(\frac{\sigma}{r}\right)^{12} - \left(\frac{\sigma}{r}\right)^{6}\right), r < r_{cut} \qquad S3$$

Here, ε= $\varepsilon_R$, the parameter determining the condensation state of the RNP, which can be 0.1, 0.2 or 0.3 kcal/mol. In the main text, the condensation states are labeled as 10* $\varepsilon_R$. The bead diameter, σ, is 1.0nm.

***RNP Model 2 (2 Components)***

RNP Model 2 includes a second type of polymer bead, S, along with the first type of polymer bead, R. The first 120 monomers are of type S (approximately the length of the 5' untranslated region (UTR) in the viral RNA), while the rest are type R. All monomers are bonded the same way as in RNP Model 1. The Lennard-Jones ε from Eq. S3 is now 3 different parameters: $\varepsilon_{RR}$, $\varepsilon_{RS}$, $\varepsilon_{ss}$. $\varepsilon_{RR}$ determines the condensation state of the majority of the RNP, while $\varepsilon_{ss}$ determines the condensation state of the signal domain. $\varepsilon_{RS}$ determines the mixing behavior between the two domains. By ensuring that $\varepsilon_{RS} \leq (\varepsilon_{RR} + \varepsilon_{SS})/2$, we ensure that the signal domain remains relatively separate from the rest of the RNP. In the main text, $\varepsilon_{RR}$ = .3 kcal/mol, $\varepsilon_{RS}$ = .2 kcal/mol, and $\varepsilon_{SS}$ = .1 kcal/mol. There is one exception when $\varepsilon_{CA\text{-}R}$ = 0 and then $\varepsilon_{SS}$ = .3 kcal/mol.

***RNP Model 3 (L-Integrase Tetramers)***

RNP Model 3 is more specific than the previous 2 models and is based on previous work modeling the RNP of HIV-1 without the capsid(*5*). Thus, different potentials are used from the previous 2 models are used to create a model that is as similar as possible to that work. RNP Model 3 consists of 2 polymers, which are each 3000 beads in length, and 35 integrase tetramers.

*RNA/NC*

Each polymer bead represents 3 bases and bound nucleocapsid protein in the dimeric RNA genome which is approximately 9000 base pairs long. Each bead has a diameter which is enforced by a soft exclusion potential shown below.

$$U_{soft}(r) = A\left[1 + \cos\left(\frac{r}{r_{cut}}\right)\right], r < r_{cut} \qquad S4$$

A is 10kcal/mol and $r_{cut}$ is 3 nm. The same harmonic potential is used to bond consecutive monomers with $r_0$ = 1.0 nm and K=5 kcal/mol/nm$^2$. We also apply these bonds to the secondary structure contacts found in the 5' UTR according to the map from Goodsell et al(*5*). This includes the contacts between the two different chains in the dimerization sequence of the RNA. We also add angle potentials

$$U_{Angle}(r) = K_\theta\ (\theta - \theta_0)^2 \qquad S5$$

To prevent unphysical bending of the RNA chains. The equilibrium angle is completely straight, 180°, although the bending constant, $K_\theta$,is a relatively weak 1.5 kcal/mol/radian$^2$.

Instead of the Lennard-Jones potential, we use a gaussian attraction to model the condensation of the RNA globule due to electrostatic interactions between NC and the RNA.

$$U_{Gauss}(r) = -\frac{H'}{\beta\sqrt{2\pi}}\exp\left(-\frac{(r-r_0)^2}{2\beta^2}\right) = -H\exp\left(-\frac{(r-r_0)^2}{2\beta^2}\right), r < r_{cut} \qquad \text{Sf6}$$

Here, $\beta$=0.5nm to give a wider range to the gaussian interaction representing the flexibility of NC and the electrostatic nature of the interaction. The equilibrium distance, $r_0$, is 3.0 nm to match the bead diameter of the repulsive potentials. It is believed that the secondary structure of the 5' UTR leads to preferential binding of NC outside of the 5' UTR in the rest of the viral RNA and thus we have different interaction strengths for beads in 5' UTR and outside- $H'_{RR}$, $H'_{UR}$, $H'_{UU}$. $H'_{RR}$ is 0.3 kcal/mol/nm, $H'_{UR}$ is 0.2 kcal/mol/nm, and $H'_{UU}$ is 0 meaning we simply apply the repulsive potential. The interactions between beads in the 5' UTR is largely given by the bonds which hold together the secondary structure.

*L-Integrase Tetramers*

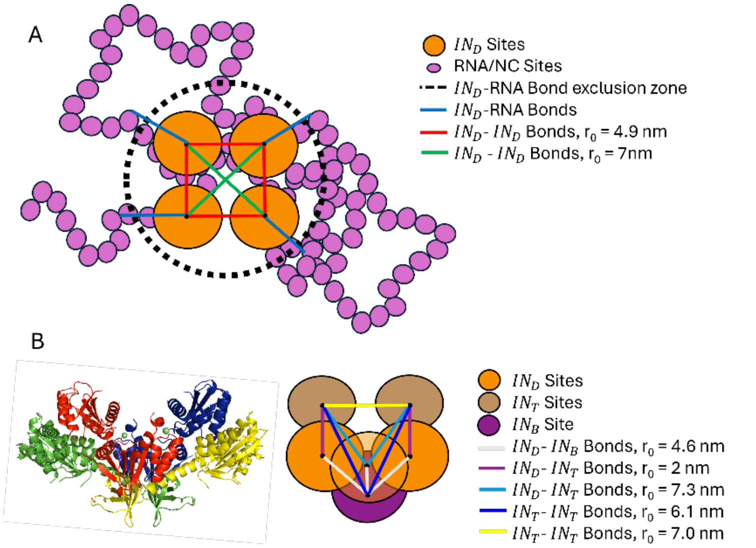


*Figure S6. (A) Model for the bonded topology of D-integrase sites. (B) The integrase tetramer structure colored by chain compared to the elastic network model used in this work.*

The integrase tetramers are based on structural data for the integrase tetramer(*6*). The integrase tetramers consist of 7 beads. They consist of 7 beads, 4 of which are arranged in a diamond like the previous IN model and are referred to as D-Integrase sites ($IN_D$). These sites, together with two sites on the top (T-Integrase) and one site on the bottom (B-integrase) form an elastic network where K is 50000 kcal/mol/nm$^2$ as shown in Figure S6. Integrase units are also bonded to the RNA/NC beads. Using the first model with integrase tetramers and self-avoiding gag from Goodsell et al.(*5*) as the initial structure, each individual integrase $IN_D$ site is bonded to the RNA/NC bead they are closest to, not including beads which are less than 7.0 nm from the center of the tetramer. This ensures that no RNA/NC bead is bound to more than integrase unit. The equilibrium distance for these bonds is set to the distance in the initial structure. Interactions between the integrase units and RNA/NC beads are given by a gaussian attraction shown in Eq. S6 in most cases. Here, $\beta$=0.5nm and the equilibrium distance, $r_0$, is 4 nm. The strength of the interaction is given by $H'_{IN-R}$ which is a parameter that ranges from 0 to 1 kcal/mol/nm to simulate any non-specific interactions between NC and integrase and to modify the amount of surface accessible integrase that CA can interact with. When $H'_{IN-R}$ is 0, the potential is replaced by a soft repulsion of the form given in Eq. S4 with $r_{cut}$=4 nm and A=10 kcal/mol.

**RNP Model 4 (Hyper-mobile Integrase)**

We remove the bonds between the Integrase monomers and the RNP polymer. We also remove the excluded volume interactions between the Integrase beads and the RNP beads so that the integrase is less impeded. The attractions between Integrase and the RNP beads (through the parameter $H_{IN-R}$) are still applied and are necessary to keep the integrase in the RNP without the bonds. $H_{IN-R}$ is always chosen to be 0.4 kcal/mol/nm as this keeps the Integrase in the RNP while allowing a similar amount of surface to integrase to what was observed previously.

**RNP Model 5 (Integrase Filaments)**

*AA Simulations of an Integrase 16 mer*

Filament Model Preparation

We used the initial all-atom PDB structure of the integrase filament bound to the CA lattice(*7*) to run an atomistic simulation as the basis for our coarse-grained model of integrase. The initial structure contained only one complete integrase octamer and small pieces of other octamers as shown in Figure S7. Each chain of the octamer was also missing many residues and the residues that were missing varied for each chain. Thus, we used MSBack(*8*) to prepare the structure. We used the chroma diffusion module(*9*) to diffuse an AF-predicted structure of the integrase monomer into each chain (8 different runs) of the octamer under the constraint that the backbone alpha carbons had to match what was given in the initial structure. We then used flowback(*10*) to predict all sidechains, added the zinc back from the initial structure and made a chain of 2 octamers by copying the octamer and aligning it to the positions of the partial octamers in the original structure. We choose to simulate a dimer of octamers since it would contain multiple copies of each interface present in an octameric filament and still be tractable to efficiently run the all-atom simulation.

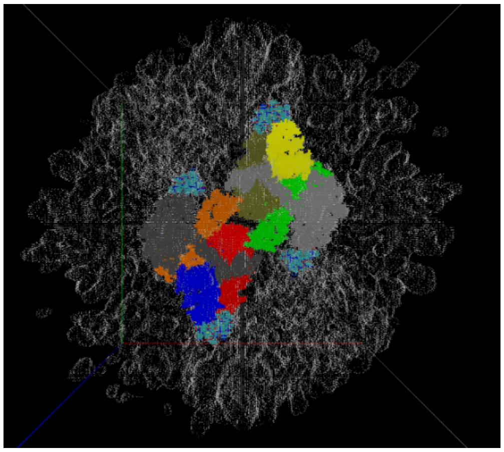

***Figure S7****. Cryo map of the integrase octamer is shown in various colors corresponding to individual chains. The capsid lattice is shown in white, and the additional pieces of integrase are shown in cyan.*

All-Atom Simulation Parameters

All all-atom MD simulations of the M protein were performed using GROMACS2022. The final 16-mer dimer of octamer structure from the methods section was used as the input. The box size was (25x25x25) $nm^3$. TIP3P water is added to the system to generate the solvated structure. We neutralized the system with a NaCl concentration of 150 mM in addition to the Zn ions present in the initial structure. The periodic boundary condition was active across the three dimensions of the simulation box. The CHARMM36 force field was used with the particle mesh Ewald sum method for long-range electrostatic interactions(*11*). After energy minimization with the steepest descent algorithm, the system was equilibrated for 20 ps in the constant NVT ensemble at 298K, followed by 100 ps at 1 atm and 298 K int the constant *NPT* ensemble. The production runs were performed for 125 ns with a 2 fs time step in the constant *NPT* ensemble using Parinello–Rahman pressure coupling and Nose-Hoover temperature coupling.

*KMC-CG + Interaction Site Mapping*

Each integrase monomer contains 273 amino acids, meaning that each tetramer contains 1092 amino acids and each octamer contains 2184 amino acids. This creates a trade off where a high-resolution model of the octamer would require many beads and thus a very costly HENM(*12*) model since the number of bonds goes as the density of beads in the protein volume O(N^3), where N is the number of CG beads. However, we also want atomistic level detail for the non-bonded interactions between the CA and the Integrase, preserving the information from the AA structure. Thus, we opted for a hybrid model where we use KMCCG mapping(*13*) to map the octamer to 120 CG beads (~18:1 mapping) but also include a 1:1 mapping of the 5 amino acids in each monomer (GLU69, TRP131, GLU138, ASN144, ASN254) that were originally suggested to interact with the CA in the AA structure for a total of 120 + 5*8 = 160 bead representation of the octamer. Excluded volume interactions will not be considered for the 1:1 mapped beads.

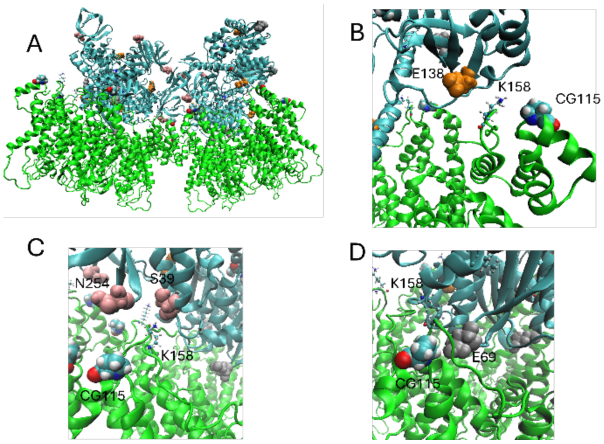


***Figure S8***. *(A) Integrase octamer is shown in cyan on top of 2 CA hexamers in green. (B-D) The proposed integrase interaction site is shown in monocolor VdW while the site of the interaction in the CG model (CG115) is colored by atom. The K158 on the CA is shown in CPK sticks. The K158 makes the all-atom contact but is not explicitly included in the CA model(1), so we choose a nearby amino acid instead.*

*HENM Inter and Intra Octamer*

A heteroelastic network was created for the integrase octamer. The all-atom simulation data was split by octamer and thus for the 125 ns of simulation we get 250 ns of training data by using each octamer separately. To create filaments of octamers of arbitrary length, we also need the elastic network parameters between the octamers. To do this, we created an henm weak elastic network where all parameters of the inter-octamer network are .1 kcal/mol/Å. This potential is not the cause of the stiffness of the filaments observed in the main text. That is mostly due to the angle potential put on the RNA which is "threaded through" the integrase filaments as discussed in the "*RNP Preparation and Relaxation*" section. In later

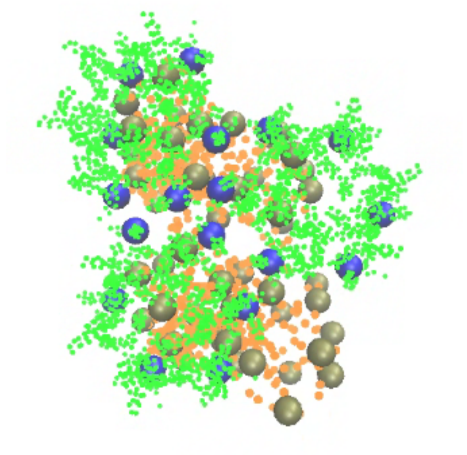

***Figure S9***. *CA hexamers are shown in green and IN tetramers in orange. Interacting sites of CA are shown in blue, while the interacting sites on IN are shown in tan.*

discussions, we realized that the filaments have a helical pitch and that our model was capturing a real phenomenon, which is impressive since it was trained on only 125 ns of a 16-mer.

*RNP Preparation and Relaxation*

Once the full model for an integrase filament of arbitrary length was established, we still needed a way to integrate it into the RNP. This was done by first building the two strands of RNA in a random walk from the dimerized 3' UTR as shown in Figure S10. The integrase filament was then positioned with the RNA strands going through the center of the filament where there are many positively charged residues. This is believed to be how the RNA binds the filament. Bonds are added between the RNA beads and all integrase beads within a 1nm cutoff such that non-bonded interactions that would normally cause clashes are excluded and the RNA is securely attached to the filament. Angle potentials are added across the section of the RNA that is within the filament to make the filament very stiff as has been described in experiments. The angles are harmonic as described by Eq. S5 with $\theta_0 = 180°$ and $K_\theta = 10000$. We can add any number of filaments of any length to the RNA polymer in this way but test only 3 filaments of length 5 and 5 filaments of length 3 in the main text due to computational limitations.

Once this configuration is created we relax the RNA polymer looking for equilibrium configurations of the RNP where integrase filaments are on the surface of the globule due to there being no attractive interactions between the filaments and the RNA while there are attractive interactions within the RNA that implicitly model the affect of positively charged nucleocapsid proteins. Relaxation is performed using molecular dynamics. First the box is compressed into a 50nm cube in 100000 CG time steps at 300k using Langevin integration. Then it is compressed into a spherical region of diameter 50 nm while the temperature is raised from 300K to 1000K over the course of 1 million CG timesteps. In the final step it is slowly cooled from 1000K to 300K over the course of 40 million CG steps. The spherical region is also increased to 70nm however the RNP does not expand due to attractive interactions. During this process we also add 5 attractive non-bonded potentials between sites on the "bottom" of the filament to make sure the integrase is correctly oriented with respect to the surface of the RNP.

*CA-IN Optimization*

We then needed to optimize the interactions between the Integrase and CA proteins. The interactions come from gaussian potentials of the form shown in Eq. S6. We started with interactions between each of the 5 sites per integrase tetramer (40 per octamer) mentioned in the mapping section and the same CA bead which has interacted with other forms of integrase but we ended up reducing to 3 sites per integrase tetramer (GLU138, GLU69, ASN254, see Figure S8). The width, β, and magnitude, H, were optimized but kept constant across the different interactions, while the minimum energy distance, $r_0$, was optimized individually for each interaction. We optimized our parameters to make sure the hexameric capsid lattice would both stably bind the filament and not be disrupted (as was the case for our initial parameter guess). We started from a 3 hexamer section of the capsid lattice bound to the integrase dimer of octamer (16-mer integrase) structure created in the Model Preparation section. We coarse-grained this configuration and iteratively ran simulations updating the mentioned parameters. H=-3 kcal/mol, β=1 Å, and $r_0$=15.5-16.6 Å, depending on the specifics on the interaction sites.

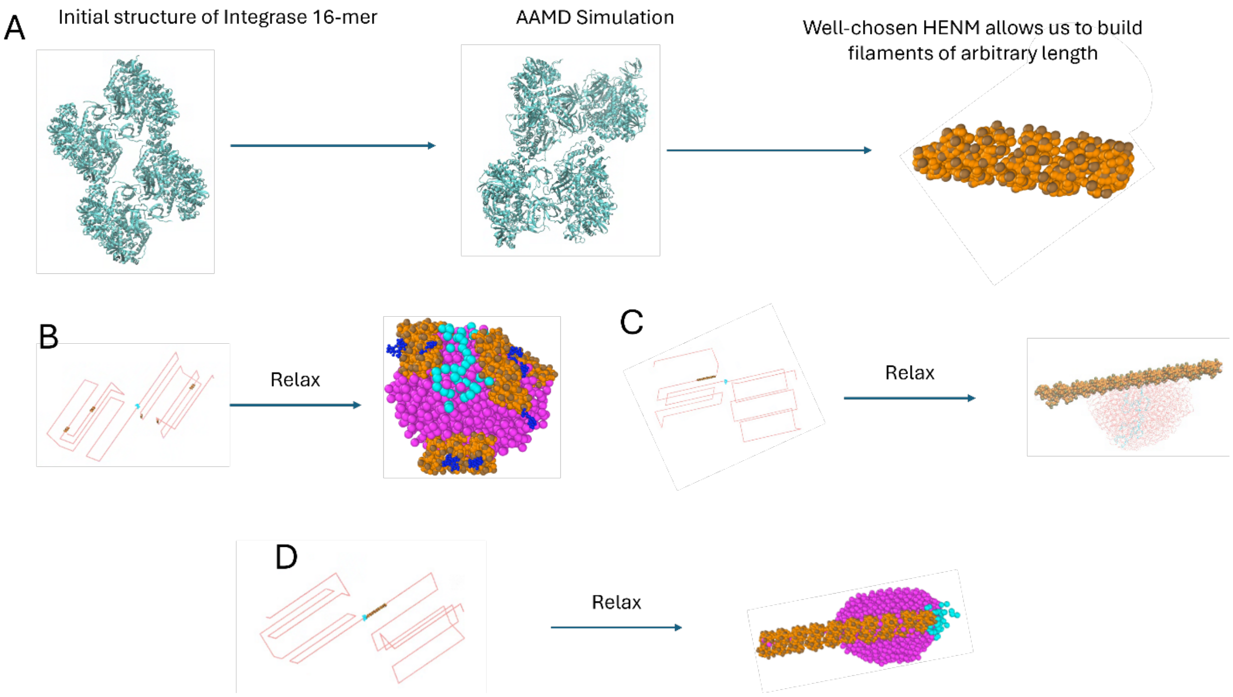


***Figure S10***. *(A) The initial model for molecular dynamics simulation of an integrase 16-mer which was prepared as described above is run for 125 ns and then the HENM is created based off of those simulation results. We can create filaments of arbitrary length as described above. (B) An example of 5 filaments of length 3 octamers being randomly placed on the RNP polymer and then relaxed to form a spherical RNP with Integrase on the surface. (CA single long filament is placed randomly on the RNP leading to the spherical RNP and the filament symmetrically on the surface, although it is longer than the RNP diameter. It becomes obvious for the long filament that there is a slight pitch angle. (D) The long filament is placed on the double stranded 5' UTR region of the RNA based on experiments suggesting the filaments interact with dsRNA. This leads it to be asymmetrically positioned on the RNP sphere.*

### RNP Model 6 (C-Integrase Tetramer)

*KMCCG + interaction site mapping, HENM*

Here, we needed a model of the integrase tetramer which would self-assemble into the octameric filaments. Thus, we opted for a hybrid model where we use KMCCG mapping to map the tetramer to 90 CG beads (~12 residue:1 CG bead mapping) but also once again include a 1:1 mapping of the 5 amino acids in each monomer (GLU69, TRP131, GLU138, ASN144, ASN254) that interact with the CA in the AA structure for a total of 90 + 5*4 = 110 bead representation of the octamer. Excluded volume interactions will not be considered for the 1:1 mapped beads. The HENM modeling was much simpler this way. We split the 125 ns of all-atom simulation data into the trajectories for 4 tetramers giving us a total of 500 ns to use to optimize the HENM.

*CA-IN and Tetramer-Tetramer Optimization*

We used the same CA-IN interactions which had previously been optimized for the tetramer including reduction from 5 interaction sites per monomer to 3. Tetramer-Tetramer interactions were chosen based on the all-atom structures and limited to a total of 10 interactions, 5 to stabilize the dimer and 5 to stabilize the filament growth. We allowed all parameters of the gaussian interactions, $r_0$ and β, to be optimized for each interaction. The parameter H' was taken as a constant for dimer and cross-mer interactions with 2 exceptions for the dimer where H* = H' - 5. This was done to prevent this interaction from over stabilizing off target states. The iterative update of parameters was carried out such that the tetrameric filament was just stable in the presence of the CA lattice but not when the CA-IN interactions were set to 0 and thus the CA lattice disassociated from the filament. The same initial condition was used as in the CA-IN optimization section for model 5. The CA-IN interactions were later further validated within the RNP for the case of hyper mobile C-Integrase in the already assembled capsid.

*Preparation of the RNP in the pre-assembled capsid*

In the case of L-Integrase this problem was trivial since the bonds just needed to be removed. In this case, we used the initial condition created by the L-Integrase assembly and removed the L-integrase, instead placing the C-Integrase tetramer at the position of each L Integrase. Since we were using the hyper-mobile version with no excluded volume interactions between integrase and the RNP polymer, there was no issue with clashes created by these configurations.

*CA Interactions with RNP Model 1 and 2*

Interactions between the RNP and CA are applied between a CG site located in the CTD of CA (CG type 115) and any bead in the RNP polymer. The potential takes the Lennard-Jones form given in Eq. S3 where $\varepsilon=\varepsilon_{CA}$ for Model 1. For Model 2 CA interacts differently with the signal domain made of S beads and thus there are two parameters, $\varepsilon_{CA\text{-}R}$ and $\varepsilon_{CA\text{-}S}$. In this paper, $\varepsilon_{CA\text{-}S}$ is always 0.9 kcal/mol for all the results while $\varepsilon_{CA\text{-}R} \leq 0.3$ kcal/mol. For both models, σ is 1.0 nm and $r_{cut}$ =2.0 nm except in the case where there are only repulsive interactions between the RNP and CA. In these cases, $r_{cut} = \sigma = 1.0$nm.

*CA Interactions with RNP Model 3*

Interactions between CA and RNP Model 3 are applied through T-Integrase ($IN_T$) sites and the same CG site as in Models 1 and 2. Justification for those choices is shown in Figure S11. All interactions between CA and the RNA/NC are given by soft repulsions as in Eq. S4, where the diameter, $r_{cut}$, is 2.0 nm. Attractive interactions come from a Gaussian potential in equation 5 where β=.5 nm, $r_0$=3nm, and H' = $H'_{CA-IN}$, the parameter which determines the attraction strength between CA and the RNP. Interactions between the other RNP beads and CA are left as a soft repulsion with a cutoff, $r_{cut}$, of 3nm.

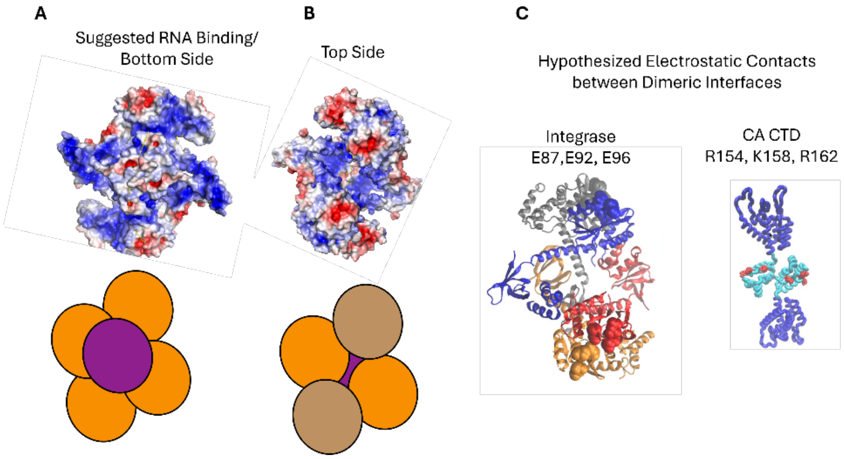


***Figure S11**. Hypothetical CA-$IN_T$ contacts used in the CG Model. (A) The positive charge (blue) on one side of the integrase tetramer suggests it binds the RNA and is not accessible to CA. (B) The other (top) side of the IN tetramer features negatively charged domains (red) on the surface where we have placed the $IN_T$ sites. (C) The integrase tetramer colored by chain shows that the 2 $IN_T$ sites are located at dimeric interfaces featuring 3 negatively charged residues per monomer (atoms shown as balls). The CA features a corresponding set of positively charged amino acids at the dimeric interface of the CA CTD where its interaction site is placed. Newer evidence suggests that double stranded RNA bound to integrase and blocked from interacting the NC may instead provide the negative charges to interact with the CA CTD. Since the model is coarse and does not explicitly contain NC, this scenario is reasonably approximated by this model.*

*CA Interactions with RNP Model 4*

Capsid-Integrase interactions are slightly adjusted such that in Equation 3, $r_0$=3.0nm instead of 2.5nm. This slight change was made after experimental results suggested the integrase may be linked to the CA through the RNA and thus it is effectively a slightly longer range interaction. This may contribute to slight shifts in the Degree of Burial distribution as it allows the integrase to be slightly more buried while still interacting with the CA.

*CA Interactions with RNP Model 5 and 6*

The optimization of these interactions were described previously in the respective RNP modeling sections.

**IP6**

IP6 anion is modeled a single bead using a center of mass mapping it interactions with all species through a soft repulsion according to Eq. S4, with one exception. That is a bead in the N-terminal of the CA monomer which can be mapped to R18, a positively charged arginine residue, 6 of which are located at the central pore of a hexamer (5 for a pentamer). That interaction is an attractive gaussian of the form shown in Eq. S6, with the parameters: β=0.04 nm, H' =-0.28 kcal/mol/nm and $r_0$ =0 .83nm. These values were used in our prior work(*2*) and calibrated based on the propensity of IP6 to bind to a pre-assembled capsid.

**Crowders**

We used an inert molecular crowding agent to represent the mass and excluded volume of Ficoll 70(*14*). Ficoll 70 was modeled using two independent spheres of CG sites connected by an elastic network of harmonic bonds with K = 1000 kcal/mol/nm$^2$. Each sphere consists of 42 CG beads and has a diameter of approximately 5.1 nm. The individual CG sites are separated by about 1 nm and a soft repulsive potential with $r_{cut}$ = 1nm is applied between the individual beads and all other species. This ensures that these spheres are impermeable, and their volume is truly excluded.

**<u>CGMD simulation details</u>**

For the CG simulations an 88 nm × 88 nm × 88 nm cubic box with periodic boundaries and with 750 CA dimers, 1700 inert crowders, 521 IP6, and one RNP is prepared. An average virion with a diameter of ~140 nm contains ~3000 CA monomers. Assuming the virion to be a perfect sphere, (volume of the virion)/(volume of the cubic simulation box with 88 nm) is ~2. Hence, we used a simulation box with half the volume of a typical virion and half as many CA (1500 CA monomer). All CGMD simulations were performed using LAMMPS (21 July 2020)(*15*). All simulations were performed in a constant NVT ensemble using a Langevin(*16*) thermostat at 300 K with a damping period of 100 ps. For nucleation simulations, a swarm of 10 parallel independent trajectories without UCG state switching were propagated for $10^7$ τ initially to expedite the identification of a nucleated cluster. The trajectory yielding a hexamer was used to initialize the production runs where UCG state switching was active. The fraction of active CA monomers ($[CA]_+$/[CA]) was 0.15, following our previous work. Experimentally observed correlation times of solution-state CA interdomain motion is 2 to 5 ns(*17*). With CG time step τ = 10 fs, state switching interval of 5 × $10^5$ τ corresponds to 5 ns of CG time. Therefore, $[CA]_+/[CA]_-$ UCG state switching was attempted every 5 × $10^5$ τ. However, caution should be exercised when comparing CG time scale to any experiments. This is a general limitation of CG methods.

*<u>Virion-like environment</u>*

For the virion-like environment, a virion-like boundary condition is created by placing particles on the surface of a 144 nm sphere to match the approximate size of the virion. The area per particle is 12 $nm^2$ /particle and they interact uniformly with all other species at an $r_{cut}$ of 4.4 nm and A=1000 kcal/mol using a soft potential as in Eq. S4 such that all particles are confined to the sphere. This doubles the volume and as such all species are doubled except for the RNP. The increased number of capsid monomers increases assembly rates and thus the fraction of active CA monomers ($[CA]_+/[CA]$) was decreased to 0.10 to observe proper assembly. This value is more in line with experimental observations in vitro(*17*).

**Data analysis and visualization**

*Definitions of hexamers and pentamers*

Graph analysis of assembled lattices was performed using the Python package NetworkX 2.1 (http://networkx.github.io/). We create a graph of CA monomers with two different types of edges, N-terminal domain edges and C-terminal domains. NTD edges are defined by the NTD dimer interface, i.e., when the distance between CG types 132/31 or 133/33 was less than 2.0 nm. CTD edges are defined between each pair of monomers in a dimer. By considering only the NTD edges we get small subgraphs that represent the oligomers present in the lattice. When these subgraphs have 5 nodes and those nodes have 10 neighbors total, that is considered a pentamer. Most hexamers are subgraphs with 6 nodes and 12 neighbors. However, we observe some hexamers where a free CA monomer has made one loose contact with a monomer in the hexamer. Thus, some hexamers have 7 nodes and 5 of those nodes have 2 neighbors, 1 node has 3 neighbors, and 1 node has only 1 neighbor. All other subgraphs are defined as disordered CA.

*Definition of Bound IP6*

In the color-coded visualizations of the capsid only bound IP6 is shown. Once the hexamers and pentamers are defined, IP6 molecules are considered bound if they are within 1.5 nm of all CG type 2 sites within a hexamer or pentamer.

*Definitions of attached CA monomers/ Interacting Integrase*

In the color-coded visualizations, CA which is not part of the growing CA lattice is still shown when it is attached to the capsid. For models 1 and 2, a CA monomer is considered attached to the RNP if CG bead type 115 is within 1.5 nm of any bead that makes up the RNP. CG bead type 115 is also the bead type that interacts with the RNP in the simulations. For model 3 with explicit integrase, CA monomers are only considered attached when CG bead type 115 is within 3.1 nm of a T-Integrase. These T-Integrase sites are also considered to be interacting.

*Estimation of Interior capsid volumes and empty space*

Using the Python package (https://github.com/scipy/scipy), capsid volumes are defined by first taking the centroids of all hexamers and pentamers. The volume of the capsid is defined as the convex hull of those points subtracted by the thickness of the capsid (1 nm) * the surface of the mesh created by the same set of points. The thickness of the capsid is a fitting parameter such that the volume capsid assembled around the most condensed one component, 18000 length polymer (18000-3) is 1.5 times the volume of the most condensed one component, 12000 length polymer (12000-3). We take these volumes to be the reference volumes for the different RNP lengths. Our assumption here is that both capsids are completely full, which is fairly accurate based on visualizations. The fraction of empty interior capsid volume in Figure 3 using model 2 is one minus the ratio of the corresponding reference RNP volume to the volume of the capsid assembled (1 – RNP reference volume/ capsid volume). For the empty fraction of the capsid interior using the explicit integrase model in Figure S3, we must define the volume of the RNP differently due to the lack of a reference state. Instead, we inspect the distance across the broad end of the capsid and subtract 2 nm for the capsid thickness based on our earlier fitting. We take half this distance to the be the radius and define the volume by assuming the RNP is a sphere.

**Definition of Clustered L-Integrase**

Clustered integrase tetramers should be those where it is not clear to the eye which tetramer an IN subunit ($IN_D$ Site) belongs to. Thus, we use a distance cutoff 6nm, which is in average of the distance between adjacent IN subunits (4.9nm) in a tetramer and non-adjacent subunits(7nm). If any $IN_D$ sites of two IN tetramers are within the cutoff then the tetramers are considered clustered.

**Definitions of Clustered C-Integrase and Integrase Filaments**

We define two different types of clusters for the C-Integrase. The first, referred to as a filament, defines a filament as a group of integrase where all tetramers are connected through the non-bonded interactions which stabilize the filaments, although there is no checking whether the structure is an octameric filament. A filament of size 6 most often corresponds to an octameric filament of length 3 as an example. The second definition, referred to as a cluster, is analogous to the definition from the L-Integrase where the intent is to say that these tetramers are physically close together. Thus, we define any two tetramers having beads less than 2nm as being in the same cluster. This cutoff distance is the same as what is used for the filaments and thus any two tetramers together in a filament are also in the same cluster.
